# Disorder in coiled-coils orchestrate chromatin organization and define a new permeability barrier at the nuclear pore basket

**DOI:** 10.1101/2025.07.18.665545

**Authors:** Zoé Narat, Stefany Figueroa, Axelle Donjon, Christine Doucet, Nouhaila Laabas, Charlene Boumendil

## Abstract

Nuclear pore complexes are the only gateway between the nucleus and cytoplasm in eukaryotic cells during interphase. Specificity of transport is ensured by a size-dependent permeability barrier, which has so far been attributed to intrinsically disordered phenylalanine-glycine (FG) repeat- containing nucleoporins located in the central channel of the nuclear pores. We show that the coiled-coil domain of the main component of the nuclear pore basket, the non-FG protein TPR, encompasses intrinsically disordered regions. This “disordered coiled-coil domain” allows it to form biomolecular condensates *in vivo*, drives mechanical exclusion of heterochromatin, and defines a new permeability barrier at the nuclear basket. This work therefore highlights unconventional properties of coiled-coil domains at the nuclear pore basket and bridges the biophysics of such domains with nuclear architecture, chromatin spatial organization and nucleocytoplasmic transport.

## INTRODUCTION

Nuclear pore complexes (NPCs) are macromolecular protein assemblies formed by multiple copies of around 35 different proteins named nucleoporins. They are embedded in the nuclear envelope through transmembrane nucleoporins that anchor their core scaffold -an inner ring sandwiched between two outer rings- forming a channel spanning the nuclear envelope. This scaffold anchors phenylalanine-glycine (FG) repeat-containing nucleoporins, which fill the central channel of NPCs. The cytoplasmic and nuclear outer rings dock the NPC cytoplasmic filaments and nuclear basket respectively^1,2^.

The NPC’s primary role is to ensure regulated transport of macromolecules between the nucleus and cytoplasm, a function that is essential for cellular physiology in all eukaryotic cells. While small molecules can freely diffuse through NPCs, larger cargoes are actively transported in a transport receptor-dependent manner. This size-dependent permeability barrier has so far been attributed to the FG repeat-containing nucleoporins located in the nuclear pore central channel^3–6^. FG repeat domains, found in central channel nucleoporins, such as NUP98, or in some peripheral nucleoporins, such as the basket nucleoporin NUP153 are intrinsically disordered regions (IDRs). IDRs do not adopt a fixed three-dimensional structure but rather sample a diverse range of conformations, enabling them to engage in transient, multivalent and weak interactions^7^. They also promote the formation of biomolecular condensates -membraneless compartments formed through multivalent interactions- which notably enable the spatial sequestration of cellular functions by concentrating specific proteins and RNAs^8^. While the exact physical properties underlying the permeability barrier of the NPCs are still under debate, the various models proposed so far -including for example the hydrogel model or dynamic polymer brush model- all agree on a key role for the molecular dynamics provided by the intrinsically disordered and cohesive nature of FG nucleoporins^1^.

In contrast with the extensively studied role of central channel nucleoporins in regulating transport, the functions and physical properties of the nuclear basket remain relatively enigmatic. The nuclear pore basket is composed primarily of the nucleoporin Translocated Promoter Region (TPR) which form long filaments through the assembly of coiled-coil homodimers^9^. Other components of the nuclear basket include the FG-repeat nucleoporins NUP153 and NUP50 and the more recently identified protein ZC3HC1, which is proposed to allow extension of TPR filaments towards the nuclear interior^10,11^. Recent advances in the structure of the nuclear basket revealed inter- and intra-species variability as well as a highly dynamic architecture^12,13^, therefore challenging the classical view of the nuclear basket as a stable and invariable structure. The basket has been implicated in various functions including mRNA export and chromatin organization, but the molecular mechanisms underlying these functions remain elusive^14–17^.

In particular, TPR has been shown to be essential for the maintenance of heterochromatin-free zones in the vicinity of the nuclear pores^15^ (Fig.1A). Indeed, heterochromatin, the transcriptionally inactive form of chromatin, is typically enriched at the nuclear periphery through its association with the nuclear lamina. Yet, in a striking contrast, the vicinity of nuclear pores is devoid of heterochromatin association. These so-called heterochromatin exclusion zones have been proposed to be necessary for proper nucleocytoplasmic transport^15^. Furthermore, our previous work demonstrated that increased NPC density during oncogene-induced senescence drives global chromatin reorganization through a TPR-dependent mechanism, implicating NPCs as active regulators of nuclear architecture, not only at the nuclear periphery but in the entire nucleus^16^. This TPR-driven chromatin reorganization is necessary for the expression of genes underlying the senescence associated secretory phenotype, therefore underlining the importance of TPR-mediated regulation of chromatin organization for the regulation of cellular state^16^. Yet how TPR maintains heterochromatin exclusion, has remained unresolved.

**Figure 1.**
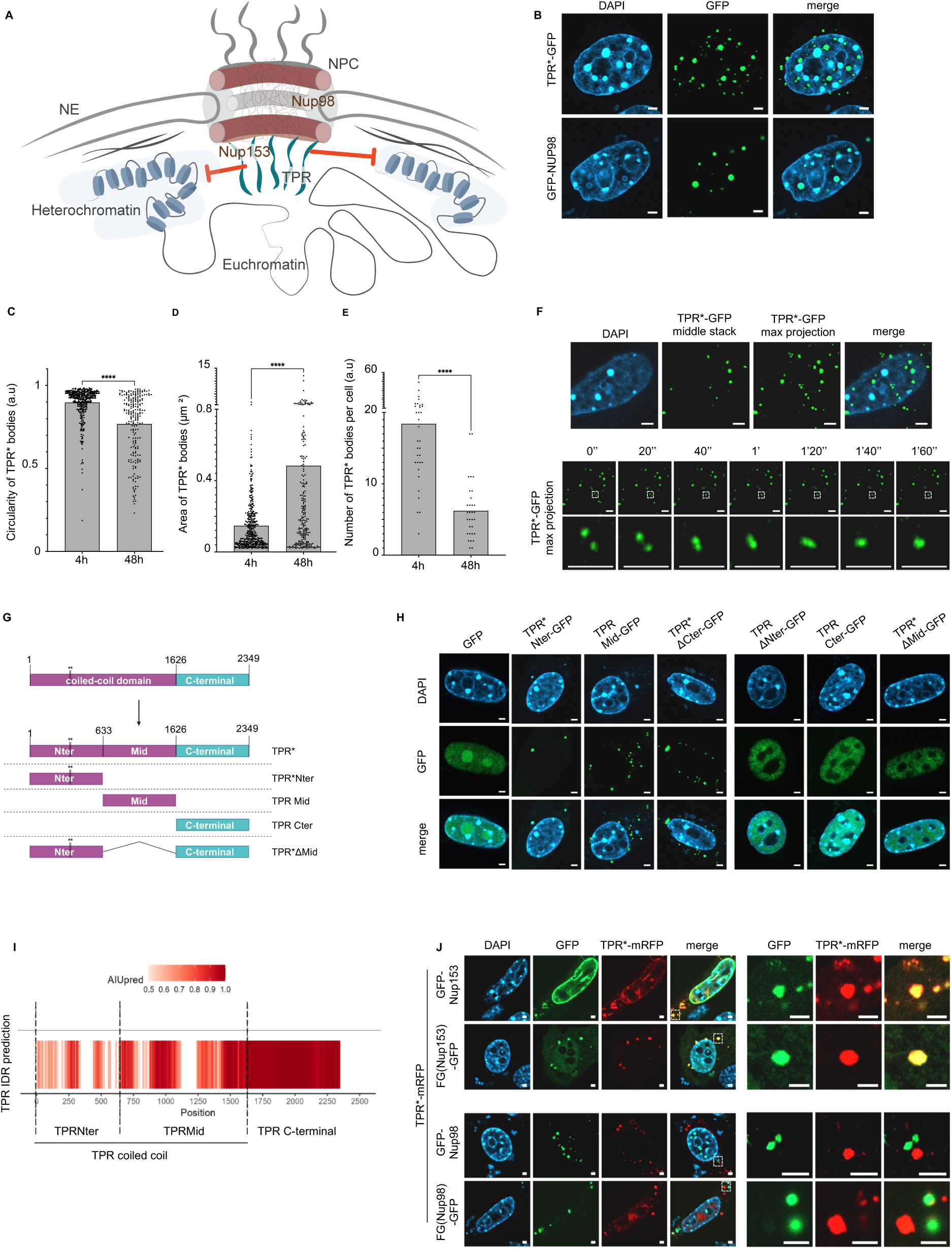
TPR’s intrinsically disordered coiled-coil drives biomolecular condensation. (A) Scheme of the nuclear pore complex (NPC) showing the position of NUP98 in the central channel, and NUP153 and TPR in the nuclear pore basket. TPR prevents heterochromatin accumulation at NPCs (red arrows). NE=Nuclear envelope. (B) Airyscan pictures of DAPI (blue), TPR*-GFP or GFP-NUP98 (green) 24h after transfection in NIH3T3 cells. A single stack is shown here. (C) Graphs showing circularity, (D) area and (E) number/cell of TPR*-GFP bodies, in stable NIH3T3 cell lines inducibly expressing TPR*-GFP, 4h or 48h upon doxycyclin induction. Each point corresponds to one body for circularity and area measurements or to one cell for number of bodies/cell. One replicate is shown here. A minimum of 30 cells has been analyzed per condition. Significance was calculated with Mann-Whitney test (****p<0.0001). (F) Top: Airyscan pictures of TPR*-GFP expressing NIH3T3 cells, 4h upon doxycycline induction (DAPI: blue, TPR*-GFP:green). Bottom: Time lapse pictures of TPR*-GFP expressing NIH3T3 cells, 4h upon doxycycline induction. Bottom panel shows a fusion event between two TPR*-GFP condensates. (G) Schematic of TPR* domains and deletion mutants. (H) Airyscan pictures of DAPI (blue) and GFP or TPR* deletion mutants-GFP (green) 24h after transfection in NIH3T3 cells. A single stack is shown here. (I) Prediction of IDRs in the human TPR protein sequence. The heatmap displays the predicted disordered propensity along the human TPR protein sequence. AIUPred colour intensity corresponds to increasing probability of a residue being part of a disordered region. (J) Airyscan picture of DAPI (blue), TPR*-mRFP (red) and GFP-NUP153, GFP-NUP98, FG(NUP153)-GFP or FG(NUP98)-GFP (green) 24h after their transfection in NIH3T3 cells. A single stack is shown here. Right panel shows a zoom on one TPR*-mRFP condensate for each condition.

## RESULTS

### Disordered coiled-coil domain of TPR drives biomolecular condensation

In order to uncover the mechanisms underlying TPR-mediated regulation of chromatin organization we set out to study TPR properties, independently of its role as a nuclear pore protein. We therefore took advantage of a TPR point mutant (TPR*), which contains two amino acid substitutions (L458D and M489D) that severely impairs its binding to nuclear pores^18^. We transfected a TPR*-GFP construct in NIH3T3 cells and surprisingly observed that rather than showing a homogenous localization, TPR* formed multiple circular bodies both in the nucleus and cytoplasm of the cells (Fig. 1B, list of all constructs generated in this study is shown in table S1). Such bodies were reminiscent of those observed upon expression of condensate-prone FG repeat-containing GFP-NUP98 fusion protein^19^ (Fig. 1B), therefore leading us to interrogate whether TPR -a non-FG nucleoporin- may be able to form biomolecular condensates. We transfected a doxycycline-inducible TPR*-GFP construct in NIH3T3 cells and followed the dynamics of TPR*-GFP bodies upon doxycycline addition. In agreement with properties of biomolecular condensates, we observed that they are circular, that their size increased over 48h while their number decreased, demonstrating coarsening of TPR*-GFP bodies over time (Fig. 1C-E and replicates in fig. S1A). Live-cell imaging of TPR*-GFP bodies 4h upon doxycycline induction showed numerous fusion and fission events, further confirming that TPR*-GFP can form biomolecular condensates in vivo (Fig. 1F, fig. S1B, Movies S1 and S2).

TPR is a 2348 amino acid-long protein, formed by a long coiled-coil N-terminal domain and an acidic, unstructured C-terminal domain ^20,21^. We therefore expected that the formation of TPR*-GFP biomolecular condensates would be led by its C-terminal domain. To address this hypothesis, we transfected several TPR*-GFP deletion mutants in NIH3T3 cells (Fig. 1G). Surprisingly however, the disordered C-terminal domain of TPR (TPR 1627-2349aa) did not form biomolecular condensates but rather showed a relatively homogenous nuclear staining (Fig. 1H). In contrast, we observed that TPR*Nter-GFP (first half of the coiled-coil domain: TPR 1-632), TPRMid-GFP (second half of the coiled-coil domain: TPR 633-1626 aa)) and TPR*ΔCter-GFP (TPR lacking its C-terminal domain: TPR 1-1626aa) formed bodies similar to those observed upon expression of TPR*-GFP or GFP-NUP98 fusion protein (Fig. 1B, Fig. 1H). Interestingly, although the full length TPR* protein is able to form biomolecular condensates, the presence of the C-terminal domain of TPR impaired the ability of TPR*Nter or TPRMid to assemble into these structures. This therefore suggests that the disordered C-terminal domain of TPR is not only unable to form biomolecular condensates but actually inhibits their formation. Presence of both Nter and Mid domains together can overcome this inhibitory role therefore suggesting an additive effect of the two domains in the formation of TPR*-GFP biomolecular condensates (Fig. 1B, 1H).

While the formation of biomolecular condensates is thought to emerge mainly from multivalent interactions between IDRs^8^, both the Nter and Mid domains of TPR are known to form α-helical coiled-coils which are the basis of the filaments structure of the nuclear basket. To understand this apparent discrepancy between the ability of these domains to form structured coiled-coils while also triggering the formation of biomolecular condensates, we applied the AIUPred prediction tool^22^ to identify potential IDRs spread out along TPR’s sequence (Fig. 1I). In agreement with a previous structural study^18^, TPR’s C-terminal domain is predicted to be fully disordered. Interestingly however, its coiled-coil N-ter and Mid domains also appeared interspersed with multiple predicted IDRs (Fig. 1I). Such coiled-coil domains interspersed with IDRs, that we thereafter refer to as disordered coiled-coils, have been previously reported in various proteins such as centrosomal proteins and were suggested to provide structural flexibility^23^. In contrast with the definition of classical IDRs, we propose that such domains may adopt a fixed 3D structure, yet sharing some physical properties of IDRs, such as cohesiveness and the ability to form biomolecular condensates.

Because various nuclear pore proteins have been shown to form biomolecular condensates through their FG-repeat IDRs^24–26^, we aimed to determine whether TPR could participate in heterotypic biomolecular condensates with FG-repeats containing nucleoporins. We therefore co-expressed a TPR*-mRFP construct and either GFP-NUP153, FG(NUP153)-GFP, GFP-NUP98 or FG(NUP98)-GFP constructs in NIH3T3 cells. As expected, overexpression of all the different constructs led to the formation of biomolecular condensates. We observed a colocalization between a fraction of TPR*-mRFP and GFP-NUP153 or FG(NUP153)-GFP biomolecular condensates, while TPR*-mRFP and GFP-NUP98 or FG(NUP98)-GFP never colocalized (Fig. 1J). Importantly, the amino acid substitutions of TPR* impair its binding to NUP153 ^18^ and the NUP153 domain known to covalently interact with TPR is located outside of its FG-repeat domain^27^, therefore eliminating the possibility of a colocalization between TPR*-mRFP and GFP-NUP153 or FG(NUP153) based on covalent interaction. These results therefore indicate that TPR* can form heterotypic biomolecular condensates with the basket nucleoporin NUP153, while not mixing with the central channel nucleoporin NUP98.

Taken together, our data show that, while not containing the canonical FG repeat motif found in condensate-prone nucleoporins, TPR can form biomolecular condensates. Surprisingly, these condensates are mediated by its coiled-coil rather than by its main disordered domain, therefore suggesting that it could have specific properties when compared to other nucleoporins.

### TPR* spatially segregates from heterochromatin domains

We next thought to determine how TPR biomolecular condensates may affect its functions. In particular, TPR has been shown to promote heterochromatin exclusion in its vicinity, both at nuclear pores^15^ and at intranuclear pools^28^, however the mechanisms underlying this function are so far unknown. We therefore aimed to test whether TPR condensates may exclude heterochromatin. To address the mechanisms of TPR - dependent heterochromatin exclusion zones maintenance independently of its role in the organization of the nuclear lamina^29^ or in transport ^20,30,31^, we again took advantage of the TPR* mutant, which is unable to bind to nuclear pores. We artificially targeted TPR*-GFP to constitutive heterochromatin, using mouse chromocentres as a model of heterochromatin domain. Chromocentres are large DAPI-dense intra-nuclear heterochromatin domains formed by the clustering of pericentric heterochromatin, which in mouse is mainly constituted by major satellite sequences (MajSat) ^32^.

To target TPR*-GFP to chromocentres, we developed inducible NIH3T3 cell lines that co-express a doxycycline-inducible catalytically dead Cas9 (dCas9) fused to a GFP-directed nanobody (GBP) and mRFP (GBP-dCas9-mRFP) ^33^ with a guide RNA (gRNA) targeting the MajSat ^34^. We named these cell lines iCRISPR-GHoST (inducible CRISPR-mediated GFP nanobody-guided Heterochromatin-Specific Targeting) (list of cell lines in Table S2). We further constructed iCRISPR-GHoST cell lines stably expressing GFP or TPR*-GFP under the control of a doxycycline-inducible promoter (Fig. 2A and fig. S2A-B). We observed recruitment of GFP or TPR*-GFP to chromocentres as early as 4h upon doxycycline induction (Fig. 2B) and did not notice any major changes in chromocentre morphology or enrichment for the repressive histone mark H3K9me3 between TPR* and GFP targeting (Fig. S2C). Importantly, staining of the endogenous TPR protein with an antibody that does not recognize TPR*-GFP ^35^ showed that TPR localization at nuclear pores was not affected by GFP or TPR*-GFP targeting at chromocentres (fig. S2D).

**Figure 2.**
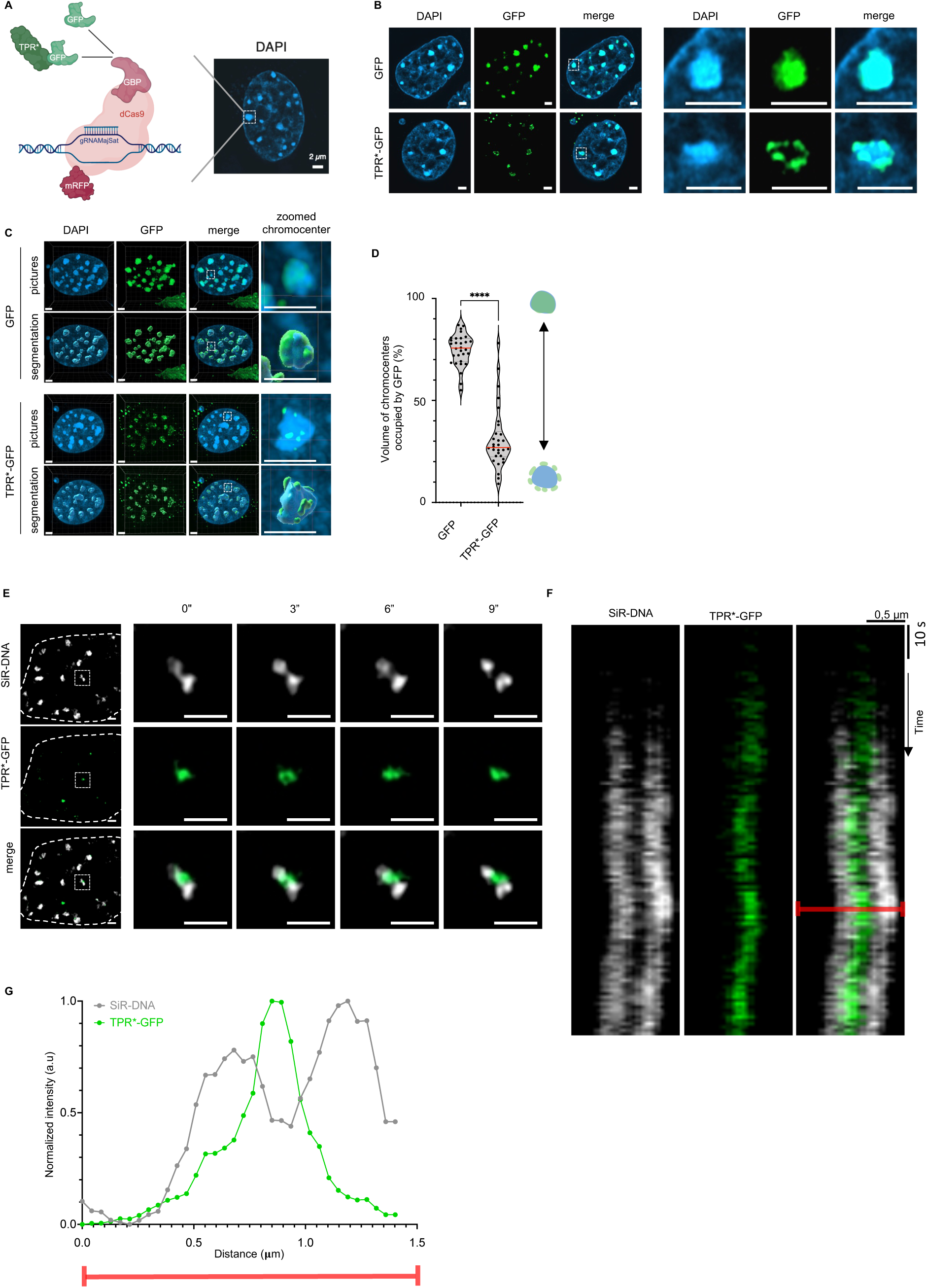
TPR spatially segregates from heterochromatin. (A) Schematic of ectopic targeting of GFP, TPR*-GFP or other proteins to chromocentres using the iCRISPR-GHOsT cell line. iCRISPR-GHOsT are NIH3T3 cells stably expressing a guide RNA against Major Satellite repeats and a GFP-binding protein (GBP)- dead Cas9 (dCas9)- mRFP fusion protein under the control of a doxycycline inducible promoter. Targeting of a protein of interest to chromocentres (here GFP or TPR*-GFP) is achieved by expressing a GFP fusion construct in these cells, either through transient transfection or stable integration of doxycycline-inducible constructs. (B) Airyscan pictures of DAPI (blue), GFP or TPR*-GFP (green) after their targeting to chromocentres in iCRISPR-GHOsT cell lines in which doxycycline-inducible GFP or TPR*-GFP constructs have been stably integrated, 4h upon doxycyclin induction. A single stack is shown here. Right panel shows a zoom on one chromocentre for each condition. (C) 3D-Segmentation of chromocentres and TPR*-GFP or GFP after their targeting to chromocentres. (D) Volume of chromocentres occupied by GFP signal (%) after targeting of GFP or TPR*-GFP in iCRISPR-GHOsT cell lines, 4h upon doxycyclin induction. A minimum of 30 cells per condition has been analyzed. Each point corresponds to one cell. The red line indicates the median. One replicate is shown here. Significance was calculated with Mann-Whitney test (****p<0.0001). (E) Time lapse pictures of DNA (SiR-DNA, grey) and TPR*-GFP (green) targeted to chromocentres in iCRISPR-GHOsT cell line, 4h upon doxycycline induction. Right panel shows a zoom on one chromocentre and highlights chromocentre rupture. (F) Kymograph generated along a line crossing the centre of a chromocentre. (G) Intensity plot of TPR*-GFP signal (green) and SiR-DNA signal (grey) across a line crossing the centre of a chromocentre at the time point indicated with a red line in panel (F).

As expected, targeting of GFP using this method led to its strong colocalization with chromocentres. In contrast, TPR*-GFP accumulated as multiple condensates at the periphery of the chromocentres (Fig. 2B). To quantify this difference, we measured the percentage of chromocentre volume occupied by GFP signal in 3D. While GFP targeting led to approximately 75% of chromocentres occupancy, TPR*-GFP only led to a 25% occupancy of chromocentre volume (Fig. 2, B-D, replicate shown in fig. S2E). To eliminate potential clonal effects, we confirmed our results in iCRISPR-GHoST transiently transfected with GFP or TPR*-GFP and included targeting of heterochromatin protein 1 (HP1) α or β as a control. We observed similar spatial segregation between TPR*-GFP and heterochromatin, while targeting of GFP or HP1 proteins led to their colocalization with the chromocentres (fig. S2 F-H). Live-cell imaging upon TPR*-GFP targeting at chromocentres further confirmed this spatial segregation and highlighted mechanical heterochromatin displacement as visualized by heterochromatin rupture upon the invasion of TPR* condensates invasion into heterochromatin domains (Movie S3, Fig.2E-G, Fig.S2I).

Interestingly, NUP98 condensates have recently been shown to be immiscible with HP1 condensates *in vitro* ^36^, therefore raising the question of whether a similar immiscibility between TPR* condensates and heterochromatin proteins could underlie the heterochromatin exclusion at nuclear pores. To test whether the presence of HP1 proteins at chromocentres was responsible for the spatial segregation with TPR*, we used previously characterized WT, HP1α KO and HP1 triple-KO bipotential mouse embryonic liver (BMEL) cells^37^ and targeted either GFP or TPR*-GFP to their chromocentres by co-transfecting them with the gRNA MajSat and GBP-dCas9-mRFP constructs. We observed that TPR*-GFP spatially segregated with chromocentres in BMEL cells, regardless of HP1 presence (Fig. S3A-B). This result therefore suggests that HP1 proteins are not necessary for TPR* and heterochromatin spatial segregation.

In sum, our data are in line with previously proposed mechanical restructuring of heterochromatin domains upon artificial induction of biomolecular condensates^38^ and led us to hypothesize that TPR* can physically exclude heterochromatin through the formation of biomolecular condensates.

### TPR* disordered coiled-coil triggers spatial segregation from heterochromatin

To test this hypothesis, we sought to determine whether the disordered coiled-coil of TPR - which promotes TPR condensate formation - is responsible for this spatial segregation from heterochromatin. For this, we targeted our multiple TPR*-GFP constructs (Fig. 1B, table S1) to chromocentres. We observed that TPRMid displayed similar spatial segregation from heterochromatin as TPR* and targeting of a deletion mutant lacking the Mid domain TPRΔMid-GFP led to its colocalization with chromocenters, therefore demonstrating a key role of the TPRMid domain in the spatial segregation between TPR and heterochromatin. TPR*Nter displayed spatial segregation from heterochromatin, although to a milder extent than TPRMid (Fig. 3, A and B, and replicates shown in fig. S4A).

**Figure 3.**
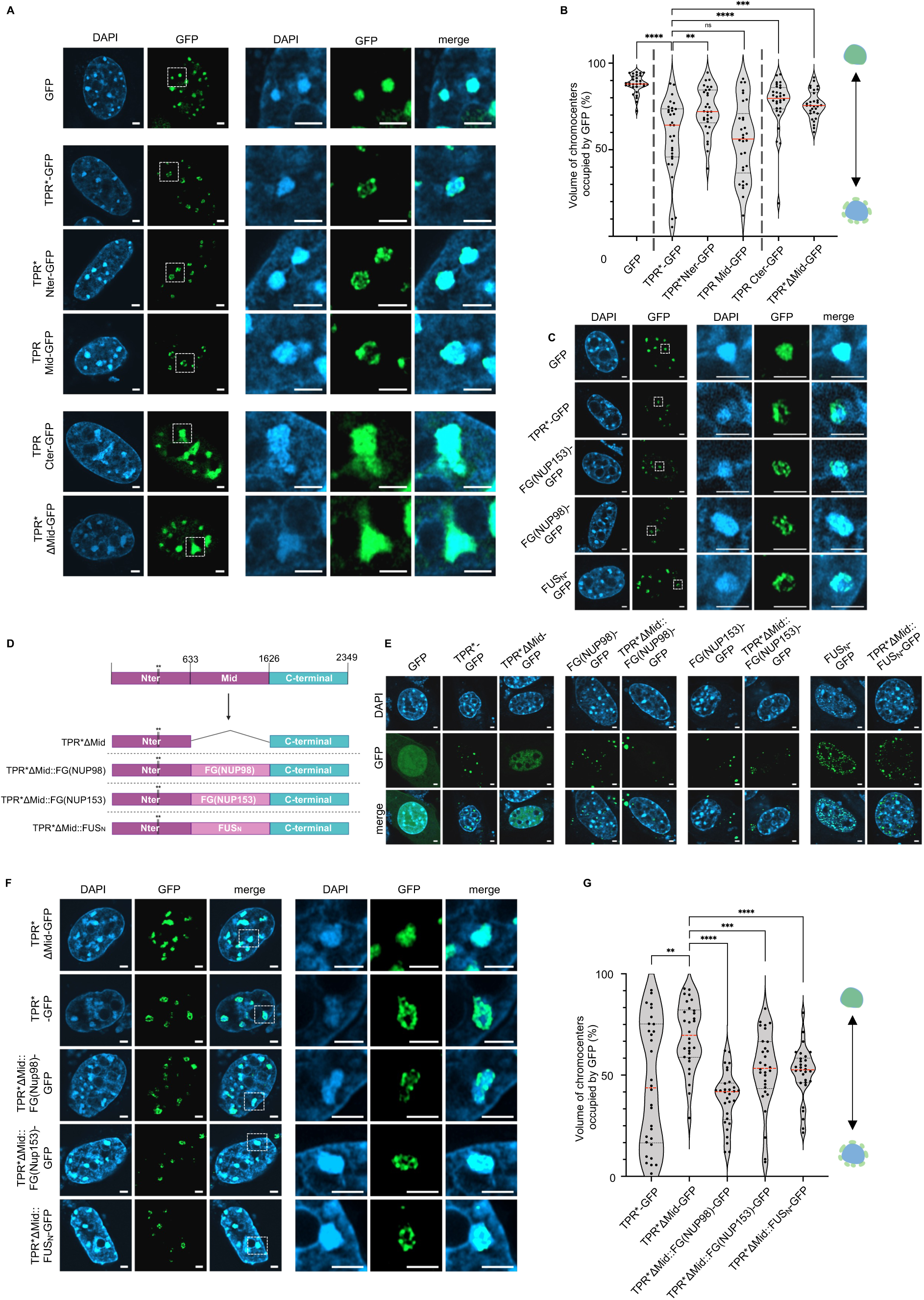
TPR biomolecular condensation is necessary and sufficient for spatial segregation from heterochromatin. (A) Airyscan pictures of DAPI (blue), GFP, TPR*-GFP, TPR*-GFP deletion mutants (green) upon their co-transfection with MajSat gRNAs in NIH3T3 cells in which a doxycycline-inducible GBP-dCas9-mRFP construct has been stably integrated, 24h post-transfection and doxycycline induction. (B) Percentage volume of chromocentres occupied by GFP signal (%), after targeting GFP, TPR*-GFP and TPR*-GFP mutants to chromocentres (as in A). A minimum of 30 cells per condition has been analyzed. Each point corresponds to one cell. Red line shows the median. One replicate is shown on this graph. Significance was calculated with Mann-Whitney test (ns: non-significant, **p<0.01 ***p<0.001, ****p<0.0001). (C) Airyscan picture of DAPI (blue), GFP, TPR*-GFP, FG(NUP153)-GFP, FG(NUP98)-GFP or FUS_N_-GFP (green) upon their co-transfection with MajSat gRNAs in NIH3T3 cells in which a doxycycline-inducible GBP-dCas9-mRFP construct has been stably integrated, 24h upon transfection and doxycyclin induction. A single stack is shown here. Right panel shows a zoom on one chromocentre for each condition. (D) Schematic of TPR* chimeras. (E) Airyscan pictures of DAPI (blue), TPR*-GFP, TPR*ΔMid-GFP, TPR*-chimeras-GFP, FG(NUP153)-GFP, FG(NUP98)-GFP or FUS_N_-GFP (green) 24h after transfection in NIH3T3 cells. A single stack is shown here. (F) Airyscan picture of DAPI (blue), TPR*-GFP or TPR* chimeras-GFP (green) after their transfection in iCRISPR-GHOsT cell lines, 24h upon transfection and doxycyclin induction. A single stack is shown here. Right panel shows a zoom on one chromocentre for each condition. (G) Percentage volume of chromocentres occupied by GFP signal (%), after targeting TPR*-GFP or TPR* chimeras-GFP to chromocentres (as in F). A minimum of 30 cells per condition has been analyzed. Each point corresponds to one cell. Red line shows the median. One replicate is shown on this graph. Significance was calculated with Mann-Whitney test (**p<0.01 ***p<0.001, ****p<0.0001).

These results demonstrate that the spatial segregation between TPR* and heterochromatin is mediated by its coiled-coil domain (TPR*Nter and TPRMid). We therefore sought to test whether the ability of TPR* coiled coil to form condensates was responsible for the observed spatial segregation between TPR* and heterochromatin. We therefore targeted multiple condensate-prone domains to chromocentres. Similarly to TPR*-GFP, targeting FUS_N_-GFP^38^, the FG-repeats containing domain of the nucleoporin NUP153 (FG(NUP153)-GFP) and the FG-repeats containing domain of NUP98 (FG(NUP98)-GFP) all led to their spatial segregation from heterochromatin in multiple condensates at the periphery of the chromocentres (Fig. 3C). This emphasizes the correlation between the ability to form condensates and spatial segregation from heterochromatin. To assess whether TPR condensation mediated by its disordered coiled coil is sufficient to trigger the spatial segregation of TPR* from heterochromatin, we constructed a chimeric TPR*-GFP in which the TPRMid domain is replaced by either the FG repeats domain of NUP98 (TPR*ΔMid::FG(NUP98)-GFP), the FG repeats domain of NUP153 (TPR*ΔMid::FG(NUP153)-GFP) or the FUS_N_ domain (TPR*ΔMid::FUS_N_-GFP) (Fig. 3D). Replacement of the TPR*Mid domain by either of these condensate-prone domains was sufficient to rescue the ability of TPR*ΔMid -GFP to form biomolecular condensates in NIH3T3 cells (Fig. 3E) and to spatially segregate from heterochromatin upon targeting at chromocentres (Fig. 3, F and G, and replicates in fig. S4B).

Taken together, our results therefore demonstrate that TPR* disordered coiled-coil is necessary and sufficient to allow its spatial segregation from heterochromatin domains when targeted to chromocentres, most likely through TPR* biomolecular condensates-mediated mechanical exclusion of heterochromatin.

### TPR disordered coiled coil maintains heterochromatin exclusion zones at nuclear pores

To test whether TPR disordered coiled-coil is involved in maintaining heterochromatin exclusion zones at the nuclear pore basket, we constructed a stable NIH3T3 cell line inducibly expressing either TPR-GFP or a GFP-fused TPR mutant lacking its Mid domain (TPRΔMid-GFP) (Fig4A, Fig. S5A). Indeed, we reasoned that the TPRΔMid mutant would retain its ability to bind to nuclear pores (mediated by the Nter domain^20^), while its disordered-associated physical properties would be restricted by the C-terminal domain. Using high resolution microscopy, we observed that expression of TPR-GFP or TPRΔMid-GFP led to the partial replacement of endogenous TPR at nuclear pores (Fig. 4A). Expression of TPRΔMid-GFP, unlike expression of TPR-GFP, led to the invasion of heterochromatin in front of the nuclear pores, as observed by a local increase in DAPI intensity at nuclear pores and by a higher Pearson correlation coefficient between DAPI and nuclear pore intensities measured along the nuclear envelope (Fig. 4A-D, fig. S5B and replicates in fig. S5C and D). This result suggests that the presence of disordered domains within TPR’s coiled-coil may be necessary to maintain heterochromatin exclusion zones at nuclear pores. To further confirm this hypothesis, we expressed a TPR-GFP chimera in which the Mid domain is replaced by an intrinsically disordered domain. For this, we constructed a TPRΔMid::FG(NUP98) chimera since the TPR*ΔMid::FG(NUP98)-GFP showed the more pronounced spatial segregation from heterochromatin at chromocentres amongst the different chimeras tested (Fig. 3, F and G, and replicates in fig. S4B). Expression of this chimera led to its recruitment at nuclear pores, partial replacement of endogenous TPR and heterochromatin exclusion similar to the one observed upon expression of a wild type TPR-GFP construct (Fig. 4A-D, fig. S5B and replicates in fig. S5C and D). This result therefore confirms that the presence of disordered domains within TPR’s coiled-coil is sufficient to maintain heterochromatin exclusion zones at nuclear pores.

**Fig. 4.**
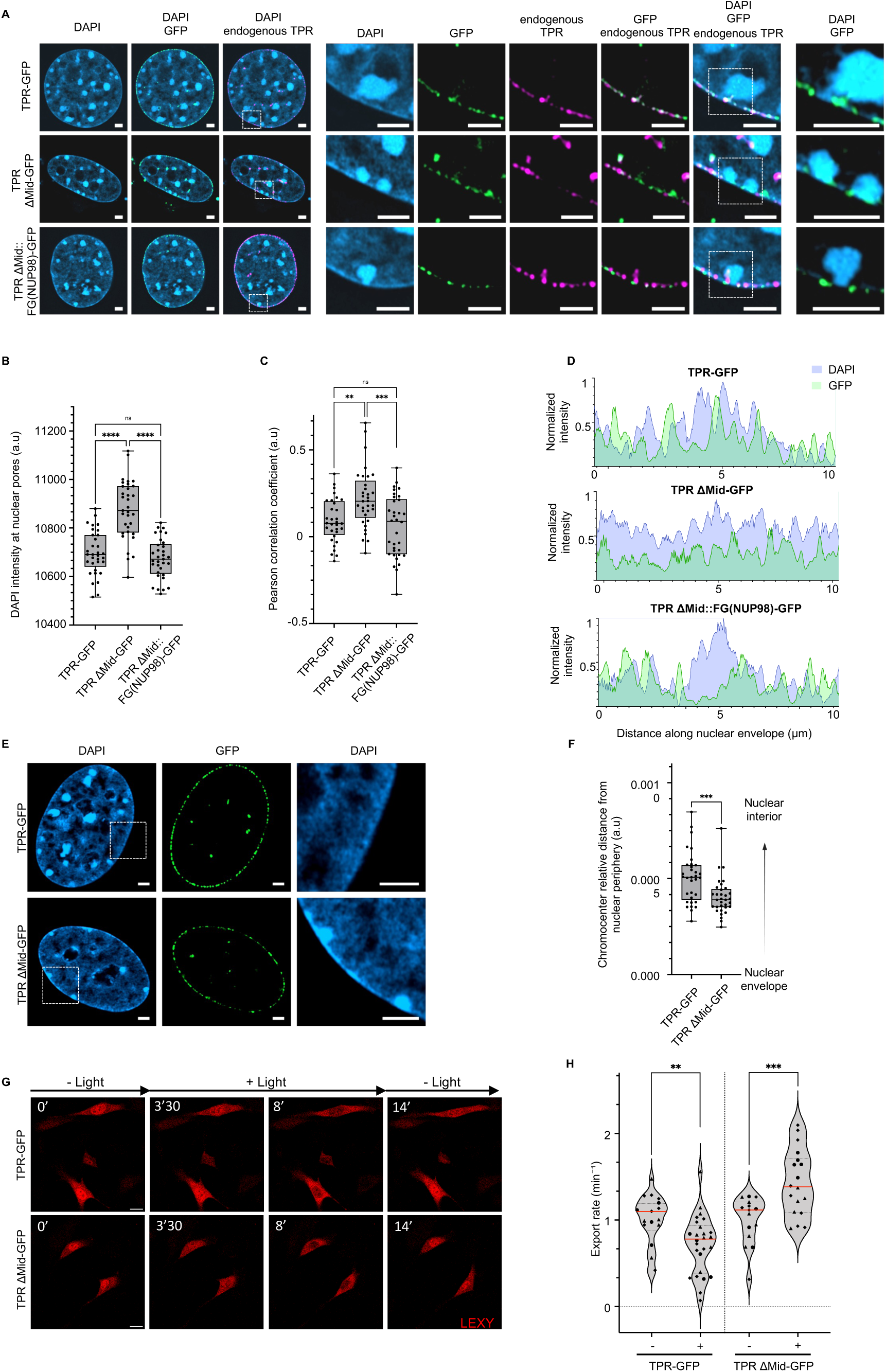
TPR’s disordered coiled coil regulates chromatin organization and nucleocytoplasmic transport. (A) Airyscan pictures of immunofluorescence staining of the endogenous TPR protein (purple) in NIH3T3 cells stably expressing either TPR-GFP, TPRΔMid-GFP or TPRΔMid::FG(Nup98)-GFP chimera (green) under the control of a doxycycline inducible promoter, 48h upon doxycycline addition. DAPI is shown in blue. (B) DAPI intensity at nuclear pores in TPR-GFP, TPRΔMid-GFP or TPRΔMid::FG(Nup98)-GFP expressing cells, as in (A). NUP153 immunofluorescence staining was used to segment nuclear pores in 3D. A minimum of 30 cells per condition has been analyzed. The interquartile range (IQR) is depicted by the box with the median represented by the center line. Whiskers maximally extend to 1.5 × IQR. One replicate is shown here. Significance was calculated with Mann-Whitney test (ns: non-significant, ****p<0.0001). (C) Pearson correlation coefficient between GFP intensity and DAPI intensity along the nuclear envelope in TPR-GFP, TPRΔMid-GFP or TPRΔMid::FG(Nup98)-GFP expressing cells, as in (A). A minimum of 30 cells per condition has been analyzed. The interquartile range (IQR) is depicted by the box with the median represented by the center line. Whiskers maximally extend to 1.5 × IQR. One replicate is shown here. Significance was calculated with Mann-Whitney test (ns: non-significant, **p<0.01,***p<0.001). (D) Intensity of DAPI and GFP signals along the nuclear envelope of one representative cell expressing TPR-GFP, TPRΔMid-GFP or TPRΔMid::FG(Nup98)-GFP, as in (A). (E) Airyscan picture of NIH3T3 cells stably expressing either TPR-GFP or TPRΔMid-GFP (green) under the control of a doxycycline inducible promoter, 48h upon doxycycline addition. DAPI is shown in blue. A single stack is shown here. (F) Chromocentres distance to the nuclear periphery normalized to nuclear volume in NIH3T3 cells expressing either TPR-GFP or TPRΔMid-GFP under the control of a doxycycline inducible promoter, 48h upon doxycycline addition as in (E). A minimum of 30 cells per condition has been analyzed. The interquartile range (IQR) is depicted by the box with the median represented by the center line. Whiskers maximally extend to 1.5 × IQR. One replicate is shown here. Significance was calculated with Mann-Whitney test (***p<0.001). (G) Representative confocal pictures of LEXY (red) time lapse in NIH3T3 cells expressing either TPR-GFP or TPRΔMid-GFP under the control of a doxycycline inducible promoter, 72h upon doxycycline addition. In absence of light activation (-Light), LEXY is mainly localized in the nucleus and light stimulation (+Light) triggers its export towards the cytoplasm. +Light time point***s*** shown here shows early and late export events. Scale bar: 20 µm. (H) Nuclear export rate of LEXY, determined by fitting LEXY’s nuclear intensity decrease to a monoexponential decay model. Individual cells of 3 independent experiments are shown as individual dots (circle: replicate 1, Triangle: replicate 2, diamond: replicate 3). Each measure was normalised by the mean of the export rate values from the corresponding -dox replicate. The significance between the -dox and +dox conditions was calculated with a T-test (**p<0.0021, ***p<0.0002). Between 15 and 26 cells were analysed per condition.

### TPR disordered coiled coil maintains global chromatin organization

Since TPR knockdown triggers major global chromatin rearrangement in senescent cells rather than a localized chromatin reorganization at the nuclear periphery ^16^, we next asked whether TPR disordered coiled-coil could affect global chromatin organization. We assessed the effect of TPR depletion on global chromatin organization in NIH3T3 cells. As expected, we observed that TPR knockdown using siRNAs in NIH3T3 cells led to the invasion of heterochromatin at nuclear pores (Fig. S6A-D). In addition to this local chromatin effect, we further observed that TPR knockdown led to a massive chromatin rearrangement, with chromocentres being relocalized towards the nuclear periphery (Fig. S7A and S6E). This result further confirms that the maintenance of heterochromatin exclusion zones at nuclear pores regulates chromatin organization in the entire nucleus^16^. We next asked whether expression of TPR lacking its Mid domain affects global chromatin organization. While expression of WT TPR did not significantly affect global chromatin organization, expression of TPR lacking its Mid domain profoundly reorganised chromatin, with a relocalization of chromocentres towards the nuclear periphery, similar to what we observed upon TPR depletion (Fig. 4, E and F, and replicates in fig. S6F). Together these data support a role for the presence of disordered domains within TPR coiled-coil in the maintenance of global chromatin organization through physical exclusion of heterochromatin from the nuclear periphery.

### TPR disordered coiled coil regulates nucleocytoplasmic transport dynamics

Having established that TPR disordered coiled coil displays some physical properties usually attributed to FG-repeats nucleoporin, we asked whether, similarly to the FG-repeats nucleoporins from the central channel, TPR disordered coiled coil could regulate nucleocytoplasmic transport dynamics. We assessed CRM1-mediated export dynamics using the light-inducible nuclear export system (LEXY)^39,40^ in TPR-GFP and TPRΔMid-GFP overexpressing cells. Briefly, the LEXY reporter assay allows to induce nuclear export of a mRFP fused reporter using light. The quantification of mRFP nuclear intensity decay upon export induction allows to calculate export rate in the different experimental conditions. Here, we used stable cells lines expressing either TPR-GFP or TPRΔMid-GFP under the control of a doxycycline-inducible promoter and compared export rates within the same cell line either in presence or absence of doxycycline to avoid potential differences due to clonal effects. We first verified that expression of TPR-GFP or TPRΔMid-GFP did not affect nuclear pore density (Fig. S7A). We observed that overexpression of TPR-GFP led to a decreased export rate while overexpression of TPRΔMid-GFP led to an increased export rate (Fig. 4G and H and Fig. S7B). This result demonstrates that TPR disordered coiled-coil domain restricts protein export rate and therefore highlights the presence of a new permeability barrier at the nuclear pore basket.

## DISCUSSION

Here, we demonstrate that the TPR coiled-coil domain harbors intrinsically disordered regions, endowing it with properties akin to classical IDR*s,* such as those in FG nucleoporin*s,* including the capacity to form biomolecular condensates in vivo. This disordered coiled-coil domain enables TPR to exclude heterochromatin at the nuclear basket and modulate global chromatin organization, while simultaneously restricting CRM1-mediated protein export. While the nucleocytoplasmic permeability barrier has traditionally been attributed to the central channel and FG-rich nucleoporins, our study challenges this paradigm by revealing TPR as a critical peripheral regulator of nuclear pore permeability. This aligns with recent high-precision tracking of cargo transport in live cells, which demonstrates that both the central channel and peripheral regions of the nuclear pore exhibit restricted diffusional environments^41^. Our results establish that TPR’s intrinsically disordered coiled-coil domain maintains this peripheral permeability barrier, showing that the nuclear basket actively contribute to selective transport.

Critically, TPR represents the first reported case of a nuclear pore component forming self-assembling molecular condensates independent of FG-repeats. While intrinsically disordered regions in ELYS and Nup35 co-assemble with Nup98 and Nup153 in vitro, they lack intrinsic phase separation properties^42^. TPR thus expands the molecular repertoire of motifs governing nuclear pore permeability. Moreover, our findings indicate that TPR condensates exist in a distinct phase from the central channel FG-nucleoporin NUP98, yet can mix with the basket nucleoporin NUP153. This highlights diverse physical properties of intrinsically disordered proteins at different nuclear pore locales, suggesting that cargoes traversing the pore may encounter multiple, mechanistically distinct permeability barriers rather than a single homogeneous gate.

Beyond permeability, our results reveal a novel function for TPR condensates: the mechanical exclusion of heterochromatin in the nucleoplasm. This demonstrates that biomolecular condensates can actively reshape their microenvironment, not merely by concentrating specific molecules, but by physically excluding others. We further show that other condensate-prone domains similarly drive heterochromatin exclusion, suggesting that this mechanism could apply to other condensates, such as transcriptional condensates and contribute to their function. These findings are particularly relevant in light of recent data showing that TPR excludes heterochromatin to repress transposons in the developing male germline^28^. Future work will determine whether endogenous TPR, highly expressed during this stage, indeed forms intranuclear condensates to mediate this repression.

At the nuclear pore, while the precise physical state of TPR remains to be elucidated, our rescue experiments with disordered domain chimeras unequivocally demonstrate that the intrinsically disordered nature of TPR’s coiled-coil domain is essential for maintaining heterochromatin exclusion. These data provide the first evidence of a physical mechanism regulating heterochromatin positioning in the nucleus, distinct from regulation of chromatin compaction states.

Taken together, our results demonstrate that intrinsic disorder within the apparently structured coiled-coil domain of TPR modulates the physical properties of the nuclear basket, coordinating nucleocytoplasmic transport and nuclear organization, two functions essential for cellular physiology. These findings not only redefine our understanding of nuclear pore permeability but also open new avenues for exploring pathological contexts, such as oncogenic translocations involving TPR’s condensate-prone Nter domain^43–45^ and viral infections targeting its condensate-inhibitory C-terminal domain^15^. Ultimately, this work prompts a fundamental question, namely to what extent do other seemingly rigid structural proteins integrate disordered regions to dynamically regulate nuclear architecture and function.

## RESOURCE AVAILABILITY

### Lead contact

Requests for further information and resources should be directed to and will be fulfilled by the lead contact, Dr Charlene Boumendil.

### Materials availability

All unique/stable reagents generated in this study are available from the lead contact with a completed materials transfer agreement.

### Data and code availability

- All original code has been deposited at [GitHub] and is publicly available at [https://gitlab.com/boumendil1/narat-figueroa-2025.git] as of the date of publication.
- Microscopy image datasets have been deposited in BioImage Archive. Accession numbers: S-BIAD2233, S-BIAD2245, S-BIAD2246, S-BIAD2247, S-BIAD2248.
- Any additional information required to reanalyze the data reported in this paper is available from the lead contact upon request.

## ACKNOWLEDGMENTS

We acknowledge the imaging facility MRI, member of the national infrastructure France-BioImaging (https://ror.org/01y7vt929), supported by the French National Research Agency (ANR-24-INBS-0005 FBI BIOGEN), and especially Marie-Pierre Blanchard, Julio Mateos-Langerak and Baptiste Monterroso for their help with microscope settings and image analyses. We are grateful to all the Boumendil and Oldfield lab members for their advice and fruitful discussions. We are grateful to Dr Valerie Doye for her insightful advice. We thank Dr Reini De Luco for hosting the group at the initiation of the project. We thank Dr Volker Cordes, Dr Maureen A. Powers, Dr Valerie Doye, Dr Giacomo Cavalli, Dr Sebastian Bultman, Dr Sophie Polo, Dr Reini De Luco for generous sharing of plasmids and reagents. We thank Sandrine Sauzet and Dr Ivana Jerkovic for experimental advice. We thank Olivier Messina for his help with Python scripts. We thank Dr Andrew Oldfield, Dr Flora Paldi, Dr Jérôme Déjardin for critical reading of the manuscript.

This work was co-funded by the European Union (ERC, NCOre 101217494). Views and opinions expressed are however, those of the author(s) only and do not necessarily reflect those of the European Union or the European Research Council. Neither the European Union nor the granting authority can be held responsible for them. Work in the CB group is also supported by the Centre National de la Recherche Scientifique (CNRS), the Institut National du Cancer (Grant INCa PLBIO23-207-2023-180), the University of Montpellier (postdoctoral grant ACT, I-SITE Excellence program under the Investissements France 2030 conducted by the HiLight Long Thematic project (PTL)- AAP25, project NAChO, project ChOICe), the Association pour la Recherche sur le Cancer (ARCDOC42023120007508), the Agence Nationale pour la recherche (ANR-23-TERC-0015-01 and ANR-21-CE12-0039).

Work in the CD group is supported by the Centre National de la Recherche Scientifique (CNRS), the national infrastructure France-BioImaging (FBI), the Agence Nationale pour la recherche (ANR-21-CE12-0023 and ANR-24-CE95-0004). Some of the illustrations were created using BioRender.

## AUTHOR CONTRIBUTIONS

Conceptualization, CB; methodology, CB, CD; formal analysis, ZN, SF, AD, CB; investigation, ZN, SF, AD, CB; visualization, ZN, SF,AD, NL, CB; funding acquisition, CB, ZN; supervision, CB; writing – original draft, ZN, SF, CB; writing – review & editing, ZN, SF, CB, AD, NL, CD

## SUPPLEMENTAL INFORMATION

**Document S1**- Table S1 to S4

**Movie S1. TPR* forms biomolecular condensates in transfected cells.** Time lapse imaging (1 image/10 seconds) 15h upon transfection of TPR*-GFP in NIH3T3 cells.

**Movie S2. TPR* forms biomolecular condensates early upon the expression of TPR*-GFP construct.** Time lapse imaging (1 image/10 seconds) of TPR*-GFP expressing NIH3T3 cells, 4h upon doxycycline induction.

**Movie S3.** Time lapse imaging (one image/second) of TPR*-GFP (green) targeted to chromocenters (marked with SiR-DNA, grey) in iCRISPR-GHOsT cells, 4h upon doxycycline induction.

## Declaration of generative AI and AI-assisted technologies in the manuscript preparation process

During the preparation of this work the authors used Emmy (Mistral AI) for proofreading. After using this tool, the authors reviewed and edited the content as needed and take full responsibility for the content of the published article.

## METHODS

### RESOURCES TABLE

| REAGENT OR RESOURCE | SOURCE | IDENTIFIER |
| --- | --- | --- |
| Antibodies |  |  |
| Anti-TPR | Abcam | ab84516 |
| Anti-Nup153 | Abcam | ab24700 |
| Anti-GFP | Merck | 11814 |
| Anti-RFP | Invitrogen | PA1-986 |
| Anti-HDAC1 | Invitrogen | PA1-860 |
| GFP booster anti-GFP AF488 | Fisher Scientific | 17343333 |
| Anti-H3K9me3 | Diagenode | C15200146 |
| Secondary antibodies |  |  |
| Anti-Rabbit AF488 | Lifetechnologies | A-21206 |
| Anti-Rabbit AF647 | Lifetechnologies | A-31573 |
| Anti-Mouse AF555 | Lifetechnologies | A-31570 |
| Anti-Rabbit HRP | Cell signaling | no. 7074 |
| Anti-Mouse HRP | Cell signaling | no. 7076 |
| Plasmids |  |  |
| GBP-dCas9-mRFP | Gift from S. Bultmann | doi:<br>10.1080/1949103<br>4.2017.1292194 |
| gRNA_MajSat | Gift from S. Polo | doi:<br>10.1038/s41467-021-22575-5 |
| GFP | This Paper |  |
| TPR*-GFP (L458D & M489D) | This Paper | cloned from<br>vector gifted by<br>V. C. Cordes<br>Doi:10.1091/mbc<br>.12.8.2433 |
| TPR*Nter-GFP | This Paper |  |
| TPR Mid-GFP | This Paper |  |
| TPR Cter-GFP | This Paper |  |
| TPRΔNter-GFP | This Paper | |
| TPRΔMid-GFP | This Paper | |
| TPR*ΔMid-GFP | This Paper | |
| TPR*ΔCter-GFP | This Paper | |
| GFP- Nup153 | Addgene | #64268 |
| GFP-Nup98 | Gift from M. A. Powers | doi:<br>10.1091/mbc.01-11-0538. |
| FUSN-GFP | This Paper | cloned from<br>vector in doi :<br>10.1016/j.cell.2018.10.057. |
| FGsNup98-GFP | This Paper |  |
| FGsNup153-GFP | This Paper |  |
| TPR*ΔMid::FG(Nup98)-GFP | This Paper | |
| TPR*ΔMid::FG(Nup153)-GFP | This Paper | |
| TPR*ΔMid::FUSN-GFP | This Paper | |
| Lenti_TetON_GBP-dCas9-mRFP | This Paper |  |
| pKLV2.3_gRNAMajSat | This Paper |  |
| Lenti_TetON_GFP | This Paper |  |
| Lenti_TetON_Piggybac_TPR*-GFP | This Paper |  |
| Lenti_TetON_TPR-GFP | This Paper |  |
| Lenti_TetON_TPRΔMid-GFP | This Paper | |
| Lenti_TetON_TPRΔMid:FGsNup98-GFP | This Paper | |
| HIV-1 gag/pol (psPAX) | Addgene | #12260 |
| VSV-G (pMD2.G) | Addgene | #12259 |
| Transposase | Gift from G. Cavalli |  |
| LEXY | Addgene | #72655 |
| Deposited data |  |  |
| FunctionsS8_TPRchromo | https://gitlab.com/boumendil1/narat-figueroa-2025.git | Script 1 |
| Script_Analysis_Table_v8 | https://gitlab.com/boumendil1/narat-figueroa-2025.git | Script 2 |
| FunctionsS10_siTPR | https://gitlab.com/boumendil1/narat-figueroa-2025.git | Script 3 |
| Script_Analysis_Table_v10 | https://gitlab.com/boumendil1/narat-figueroa-2025.git | Script 4 |
| FunctionsS12_TPRpores | https://gitlab.com/boumendil1/narat-figueroa-2025.git | Script 5 |
| Script_Analysis_Table_v12 | https://gitlab.com/boumendil1/narat-figueroa-2025.git | Script 6 |
| TPR_foci.icpx | https://gitlab.com/boumendil1/narat-figueroa-2025.git | IMARIS batch 1 |
| Fiji_macro.ijm | https://gitlab.com/boumendil1/narat-figueroa-2025.git | Macro 1 |
| PearsonCoeff_siRNA | https://gitlab.com/boumendil1/narat-figueroa-2025.git | Script 7 |
| PearsonCoeff_TPRpores | https://gitlab.com/boumendil1/narat-figueroa-2025.git | Script 8 |
| Image datasets | BioImage Archive | S-BIAD2233, S-BIAD2245, S-BIAD2246, S-BIAD2247, S-BIAD2248 |
| Experimental models: Cell lines |  |  |
| NIH3T3 | ATCC® | CRL-1658™ |
| HEK293T | ATCC® | CRL-11268™ |
| NIH3T3_GBP-dCas9-mRFP | This Paper |  |
| iCRISPR GHoST :<br>NIH3T3_GBP-dCas9-mRFP_gRNAMajSat | This Paper |  |
| NIH3T3_GBP-dCas9-mRFP_gRNAMajSat_GFP | This Paper |  |
| NIH3T3_GBP-dCas9-mRFP_gRNAMajSat_TPR*-GFP | This Paper |  |
| NIH3T3_TPR*-GFP | This Paper |  |
| NIH3T3_TPR-GFP | This Paper |  |
| NIH3T3_TPRΔMid-GFP | This Paper |  |
| NIH3T3_TPRΔMid::FG(Nup98)-GFP | This Paper |  |
| BMEL WT | Doi:10.1038/s41388-020-1177-8 | Gift from Dr Florence Cammas |
| BMEL HP1α KO | Doi:10.1038/s41388-020-1177-8 | Gift from Dr Florence Cammas |
| BMEL TKO | Doi: 10.1038/s41388-020-1177-8 | Gift from Dr Florence Cammas |

#### Cell culture, establishment of stable cells lines and transfections

All cells were maintained at 37°C with 5% CO_2_ in Dulbecco’s modified Eagle’s Medium (DMEM) GlutaMax (Gibco, no. 31966-021) supplemented with 10% tetracycline-free fetal bovine serum (FBS) (Sigma, no. F2442) and 1% of penicillin/streptomycin (Gibco, no. 15140-122) (complete medium). Cells were routinely passaged at 70-80% confluency and splitted using trypsin 0.05% (Gibco, no. 25300). Expression of doxycycline-inducible vectors was induced using 1 µg/ml of doxycycline (Sigma, no. D9891) for a minimum of 4h (specific induction time for the different experiments can be found in the figures legend). A list of all cell lines used or generated in this study can be found in Table S2. All genetically modified cell lines were selected using antibiotic selection corresponding to the antibiotic resistance cassette of the vector and clones were selected upon clonal expansion. Antibiotics concentrations used for these cell lines are 0.5 µg/ml of Puromycin (Gibco, no. A11138), 100 µg/ml of Hygromycin (Gibco, no. 10687010) and 600 µg/ml of Neomycin (Gibco, no. 10131).

Bipotential mouse embryonic liver (BMEL) cells Wild Type (WT), HP1α KO or TKO (lacking all three HP1 paralogs alpha, beta and gamma) were established by Dr Florence M. Cammas (IGH, Montpellier). Cells were cultured at 37 °C in a humidified atmosphere with 5% CO2. BMEL cells were grown in previously coated plates for 2-3h hours with 2µg/ml of Fibronectin (Sigma, no. F1141) and 20 µg/ml of Collagen (Sigma, no. 08-115) in Roswell Park Memorial Institute medium (RPMI 1640; 61870010, Gibco) supplemented with 10% tetracycline-free fetal bovine serum (FBS) (Sigma, no. F2442) and 1% of penicillin/streptomycin (Gibco, no. 15140-122), 10 μg/mL insulin (I9278, Sigma), 30 ng/mL IGFII (I8904, Sigma), and 50 ng/mL EGF (E5160, Sigma). Cells were routinely passaged at 70-80% confluency and splitted using trypsin 0.05% (Gibco, no. 25300).

Stable cell lines were generated using either lentiviral infection or PiggyBac transposon system. For lentiviral particles production, HEK293T cells were transfected using the calcium phosphate precipitation method. A DNA mix containing 5 µg of the plasmid of interest, 5 µg of the packaging plasmid psPAX2 (HIV-1 Gag/Pol), and 1 µg of the envelope plasmid pMD2.G (VSV-G) was prepared in 450 µL of sterile H₂O. Then, 450 µL of 2× HEPES-buffered saline (Sigma, no. 51558) and 125 µL of CaCl₂ 1 M (Sigma, no. 21115) were successively added. The mixture was vortexed and immediately mixed thoroughly by pipetting. The resulting calcium phosphate-DNA precipitate was added dropwise onto HEK293T cells. 18h post-transfection, the culture medium was replaced with fresh complete DMEM. Viral supernatants were collected 48 and 72 hours after transfection and used to infect NIH3T3 cells, followed by clonal selection.

For PiggyBac-mediated stable integration, cells were co-transfected with the PiggyBac vector and a helper plasmid encoding the PiggyBac transposase using Lipofectamine 2000, according to the manufacturer’s protocol (Invitrogen, no. 52887).

For plasmid transfections, 90,000 cells were plated onto coverslips placed in 6-well plates, previously coated for 1.5 hours with 2µg/ml of Fibronectin (Sigma, no. F1141) and 20 µg/ml of Collagen (Sigma, no. 08-115). All transfections in NIH3T3 cells, NIH3T3-derived stable cell lines and BMEL were performed in complete medium (DMEM or RPMI) without Penicillin/Streptomycin, using Lipofectamine 2000 (Invitrogen, no. 52887) at a 1:200 ratio, with 2 µg of total plasmid DNA diluted in Opti-MEM medium (Gibco, no. 31985). The transfection medium was replaced 15 hours post-transfection with fresh complete medium, cells were collected for further analysis 15-24h upon transfection.

For siRNA transfections, 2µl of Dharmafect 1 (Dharmacon, no. T-2001-03), 348µl of Opti-MEM and 0.72µl of 100µM siRNA were mixed and added dropwise in the cells. The transfection media were changed between 15h and 20h post-transfection with complete medium, cells were collected for further analysis 72h upon transfection. The siRNAs used in this study are : ON-TARGETplus Non-targeting Pool (Dharmacon, reference: D001810-05), and ON-TARGETplus Mouse Tpr Smart pool (L-041153-01-0005).

#### Cloning

All constructs were generated using standard molecular cloning techniques, including PCR amplification with Q5 High-Fidelity DNA Polymerase (NEB, no. M0491) or LongAmp® Taq DNA Polymerase (NEB, no. M0323), restriction enzyme digestion, and either recombination-based assembly NEBuilder HiFi DNA Assembly (NEB, no. E2621) or traditional restriction-ligation cloning with T4 DNA ligase (NEB, no. M0202), depending on the construct. Deletion mutants of TPR* or TPR were constructed by amplifying selected fragments of the protein coding sequence, which were then assembled into NLS-EGFP vectors using HiFi. Chimeric constructs were generated by first removing the Mid domain of TPR region from the wild-type TPR plasmid, and subsequently inserting the desired sequences in its place, using HiFi. Constructs of interest were then subcloned into lentiviral vectors for stable expression, under either a TetON-inducible promoter or a constitutive promoter. To clone small fragments such as gRNAs or to add restriction sites, complementary primers were ordered from Eurofins (via Custom DNA oligos) and annealed by incubation at 98 °C for 5 minutes followed by gradual cooling overnight on a heating block. The resulting double-stranded fragments were then ligated into the plasmid backbone using T4 DNA ligase. Plasmids after cloning were all transformed with NEB^®^ 5-alpha Competent E. coli (High Efficiency) (NEB, no. C2987) and amplified by transformation with NEB^®^ 5-alpha Competent E. coli (Subcloning Efficiency) (NEB, no. C2988). All constructs were verified by DNA digestion or PCR colony method and whole plasmid sequencing was performed by plasmidsaurus using Oxford Nanopore Technology with custom analysis and annotations. A list of all plasmids is provided in Table S1 and full plasmid sequencing is available on request. All plasmids have Ampicillin resistance gene except for the plasmids GFP-NUP153 and GFP-NUP98 which are Kanamycin resistant.

#### Western blot analysis of protein expression

Cells were lysed in Laemmli buffer (SDS 2%, glycerol 10%, 0.5M Tris pH=6.8) either directly in 6-well plates (500,000 cells, 150-250µl) after scraping, or after trypsinization, PBS wash, and resuspension (4 million cells, 200µl). Lysates were boiled for 7 minutes and protein concentration was measured using a Nanodrop or with Qubit Protein BR Assay (ThermoFisher, no. A50668). 20µg of proteins were run into NuPAGE™ 3-8 %, Tris-acetate, 1.5 mm (Invitrogen, no. EA0378) in NuPAGE™ Tris-Acetate SDS migration buffer (20X) (Novex, no. LA0041). Proteins were transferred onto Nitrocellulose/Filter Paper Sandwiches (BioRad, no. #1620215) by wet transfer using NuPAGE™ transfer buffer (20X) (Invitrogen, no. NP00061) for 1h30 at 100V. The membranes were blocked in 5% milk in Tris-buffered saline (TBS) with 0.1% Tween-20 (TBS-T) for 35 min at room temperature. Membranes were then probed with primary antibodies listed in Table S3, diluted in 5% milk. After washes (5, 10, and 15 minutes) in 0.1% TBS-T, membranes were incubated for 1 hour at room temperature with HRP-conjugated secondary antibodies (see Table S3), followed by additional washes. Immunoreactive bands were revealed using either Western Pierce™ ECL transfer substrate (Thermo Scientific, no. 321006) or SuperSignal™ West Femto Maximum Sensitivity Substrate (Thermo Scientific, no. 34090), depending on the signal intensity. Chemiluminescence was detected using a ChemiDoc™ MP Imaging System for Reliable and Reproducible Multiplex Fluorescent Imaging (Bio-Rad Laboratories, Hercules, CA, USA). Band quantification was performed with Image Lab software (Bio-Rad).

#### Immunofluorescence

For all immuno-staining experiments, all steps were performed at room temperature. Cells plated on coated coverslips (2µg/ml of Fibronectin (Sigma, no. F1141) and 20 µg/ml of Collagen (Sigma, no. 08-115)) were washed in PBS 1X and fixed with 4% of Pierce™ Formaldehyde (w/v), Methanol-free, (Thermo Scientific, no. 28908) diluted in PBS 1X. Cell fixation was followed by permeabilization in 0.5% Triton X-100 (Dutscher, no. T8787) diluted in PBS 1X for 10 minutes. Coverslips were then blocked with 1% bovine serum albumin (BSA) (Sigma, no. A3059) in PBS 1X for 30 minutes and incubated with the primary antibody (Table S4) diluted in 1% BSA for 1 hour in a humidified chamber. Cells were washed in PBS 1X three times for 3 minutes and incubated with the corresponding secondary antibody for 1 hour in a humidified chamber. Nuclei were then stained with DAPI (Sigma, no. D9542 diluted in PBS 1X at 0.2 µg/ml for 2 min and washed again in PBS 1X three times for 3 minutes. To remove salts, coverslips were finally rinsed with water (Sigma, W3500) for 5 min. All coverslips were mounted with VECTASHIELD^®^ Antifade Mounting Medium (H-1000-10).

#### Super Resolution Imaging

Imaging of fixed cells was performed using a Zeiss LSM980 confocal inverted microscope equipped with an Airyscan 2 module, operated in super-resolution mode with a 63×/1.4 oil immersion objective, the zoom factor was different for each experiment to image single cells. GFP, mRFP, and far-red fluorophores (Alexa Fluor series) were excited with 488 nm, 543 or 555 nm and 647 nm lasers, respectively. Emissions were detected using the following spectral range: 400-505nm for DAPI, 400-550nm for GFP or AF488, 525-720nm for mRFP or AF555 and 640-720nm for SiR-DNA or AF647. Airyscan lasers alignment were calibrated before each session for the 4 different channels aligning the visible channels on Live mode and the UV on Continuous mode with one of the visible channels. Full nucleus 3D imaging was acquired with the following airyscan settings: 0.2µm z-steps, dual time of 1.15 µs and a size of 0.31 µm. Airyscan image processing was applied automatically after imaging using Zeiss ZEN software (version blue 3.6) to obtain the final processed images.

#### Live cell imaging

Live-cell imaging was conducted using the same microscope in fast Airyscan mode (SR-4Y configuration), under environmental control (37 °C, 5% CO₂). For all live-cell experiments, cells were maintained in phenol red-free DMEM supplemented with 10% FBS and 2 µl of SiR-DNA 647 (Spirochrome). Doxycycline was added to the medium at the indicated timepoints. Time-lapse imaging was performed every 10 seconds for fusion-fission experiment and every second for kymograph creation upon TPR* targeting at chromocenters.

#### Nucleocytoplasmic transport assay

Live cell imaging was performed using the confocal microscope SP8 with a resonant scanner, and a 40x oil immersion objective, under environmental control (37°C, 5% C02). The assay was performed in triplicate, and the following analyses were realised as in^40^. The export rate measures were normalised by the mean of the -dox export rate from the corresponding replicate, for both cell lines.

#### Image analysis

Processed images obtained by the Airyscan were analyzed using IMARIS 10.0.0 (Bitplane, RRID:SCR_007370) with the XT module, unless otherwise specified.

To quantify the chromocentres occupancy, ie percentage of chromocentres volume occupied by GFP, chromocentres were segmented using the surface tool on the DAPI channel with a 0.05µm grain size, manual segmentation threshold and a filter discarding all the surfaces under 1µm^2^ area that we did not consider as “chromocentres”. The green signal corresponding to GFP, TPR*-GFP, or TPR*-GFP mutants targeted to the chromocentres was also segmented using the surface tool on the 488 channel with a 0.05µm grain size, manual segmentation threshold and a filter discarding all the surfaces under 0.3 µm^2^ area. The nucleus was segmented on the DAPI channel using the surface tool, grain size 1µm and manual threshold. A Distance transformation was done on the chromocentres outside to have a measure of overlapping with the GFP segmented surfaces. Colocalization volume was calculated by summing up the volume in the segmented GFP that overlapped/colocalized with the segmented DAPI, dividing it by the sum of all segmented chromocentres in DAPI, and multiplying it by 100 to have a percentage. The resulting percentage reflects the proportion of chromocentres volume occupied by GFP signal within the nucleus. Data exported from IMARIS were sorted using a custom Python script (Table S4, script 1 and 2) and plotted with GraphPad Prism (v.10.3.1).

For the analysis of chromatin organization at the nuclear periphery, IMARIS with the XT module was used, chromocentres segmentation was done as previously described, the nucleus segmentation was performed with the surface tool drawing the contours of the nucleus on different z stacks manually and then combined to obtain the whole nucleus volume. Nuclear pore complexes were segmented using the spots tool set at 150nm. A filter based on the shortest distance to the nuclear surface was applied to exclude spots located away from the nuclear membrane. With the XT module, the shortest distance between each chromocentre and the nuclear envelope was automatically calculated to quantify their positioning relative to the nuclear periphery. To obtain the DAPI intensity specifically at NPC, we extracted the DAPI intensity only inside the segmented NPC spots and calculated an average DAPI intensity for each cell. This average DAPI intensity was then normalized with the whole nucleus intensity. Data exported from IMARIS were sorted using a custom Python script (Table S4, script 3 to 6) and plotted with GraphPad Prism (v.10.3.1).

To quantify the circularity, area, and number of GFP-positive foci, the surface tool of IMARIS was used in the green channel with a 0.07µm surface grain size, coupled with machine learning-based segmentation tools previously trained to identify and analyze the individual objects. An area filter of 0.02µm-100µm and a circularity filter of 0.05µm-1µm was applied (Table S4, IMARIS batch 1).

To analyze the Pearson correlation coefficient and extract fluorescence intensity profiles along the nuclear periphery, a custom macro was developed in Fiji (V.2.16.0) (Table S4, Macro 1). For each image, one Z-plane was manually selected and extracted. The nucleus was segmented based on a modular threshold applied to the DAPI signal. To select the nucleoplasm layer just adjacent to the nuclear pores, the mask was rescaled by 2 % in both X and Y dimensions using a centered geometric transformation. Fluorescence intensity profiles were then measured along the eroded segmentation border on each channel. Pearson correlation coefficient was calculated by a python script (Table S4, script 7 and 8) and data were plotted with GraphPad Prism.

Kymographs were generated using the Fiji plugin “KymographBuilder”, with a line drawn across one chromocenter. Quantification of TPR*-GFP (green) and chromocenter (SiR-DNA, far red) intensity has been performed by drawing a horizontal thick line on the kymograph. Intensity has been normalized to min and max (0-1) and plotted with GraphPad Prism.

#### IDRs prediction

Intrinsically Disordered Regions (IDRs) were predicted using IUPred3 with ‘only disorder’ and ‘default smoothing’ options enabled^22^. Human TPR protein sequence was retrieved from the UniProt database, accession P12270. TPR IDRs heatmap was plotted using R package ‘ggplot2’ version 4.0.1. A threshold of 0.5 was applied to plot residues with more than a 50% probability of being part of a disordered region.

