## Supplementary figures and tables for "Disorder in coiled-coils orchestrate chromatin organization and define a new permeability barrier at the nuclear pore basket"

**The PDF file includes:**

Figure S1 to S7

Legends of Movies S1 to S3

Table S1 to S4

**Other supplementary materials for this manuscript include the following:**

Movies S1 to S3

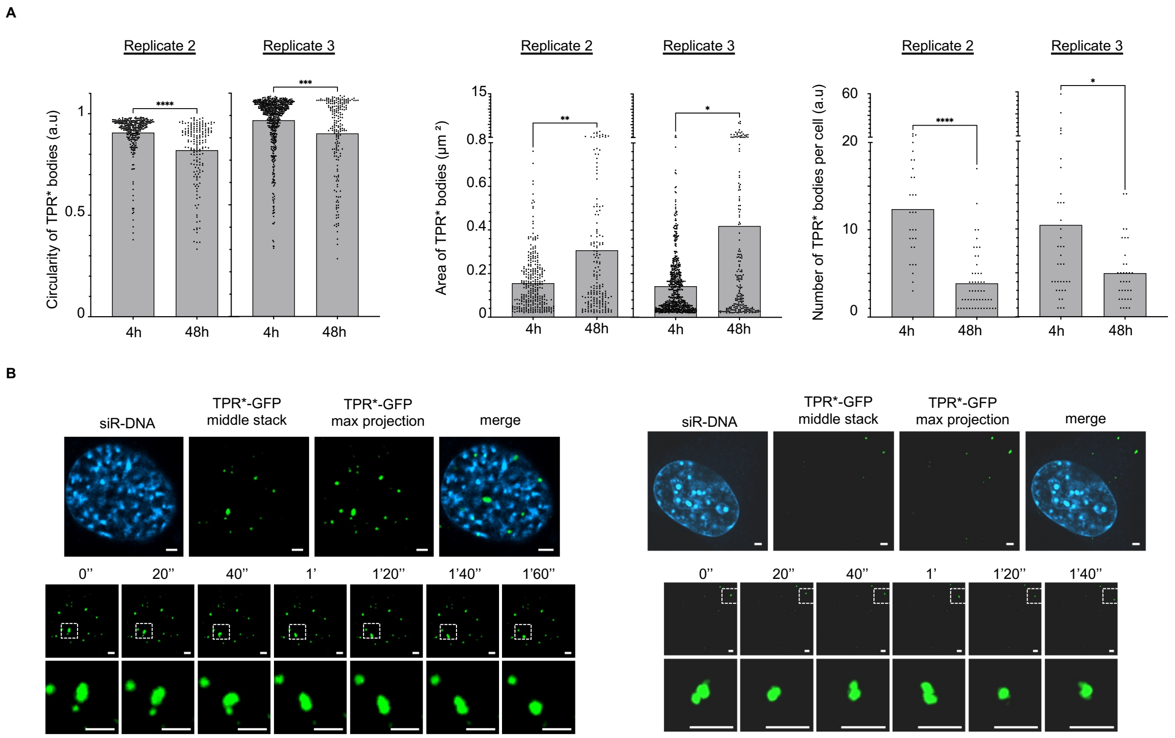
A
Replicate 2
Replicate 3
Circularity of TPR* bodies (a.u)
Area of TPR* bodies (μm ²)
Number of TPR* bodies per cell (a.u)
4h
48h
B
siR-DNA
TPR*-GFP middle stack
TPR*-GFP max projection
merge
0''
20''
40''
1'
1'20''
1'40''
1'60''

Figure. S1.

**Fig. S1. TPR* forms biomolecular condensates.**

(A) Graphs showing circularity, area and number/cell of TPR*-GFP bodies in stable NIH3T3 cell lines inducibly expressing TPR*-GFP, 4h or 48h upon doxycyclin induction. Two independent biological replicates are shown here. Each point corresponds to one body for circularity and area measurements or to one cell for number of bodies/cell. A minimum of 30 cells has been analyzed per condition. Significance was calculated with Mann-Whitney test (*p<0.05, **p<0.01, ***p<0.001, ****p<0.0001). (B) Time lapse pictures of TPR*-GFP expressing NIH3T3 cells, 4h upon doxycycline induction. Bottom panels show fusion and fission events between TPR*-GFP condensates.

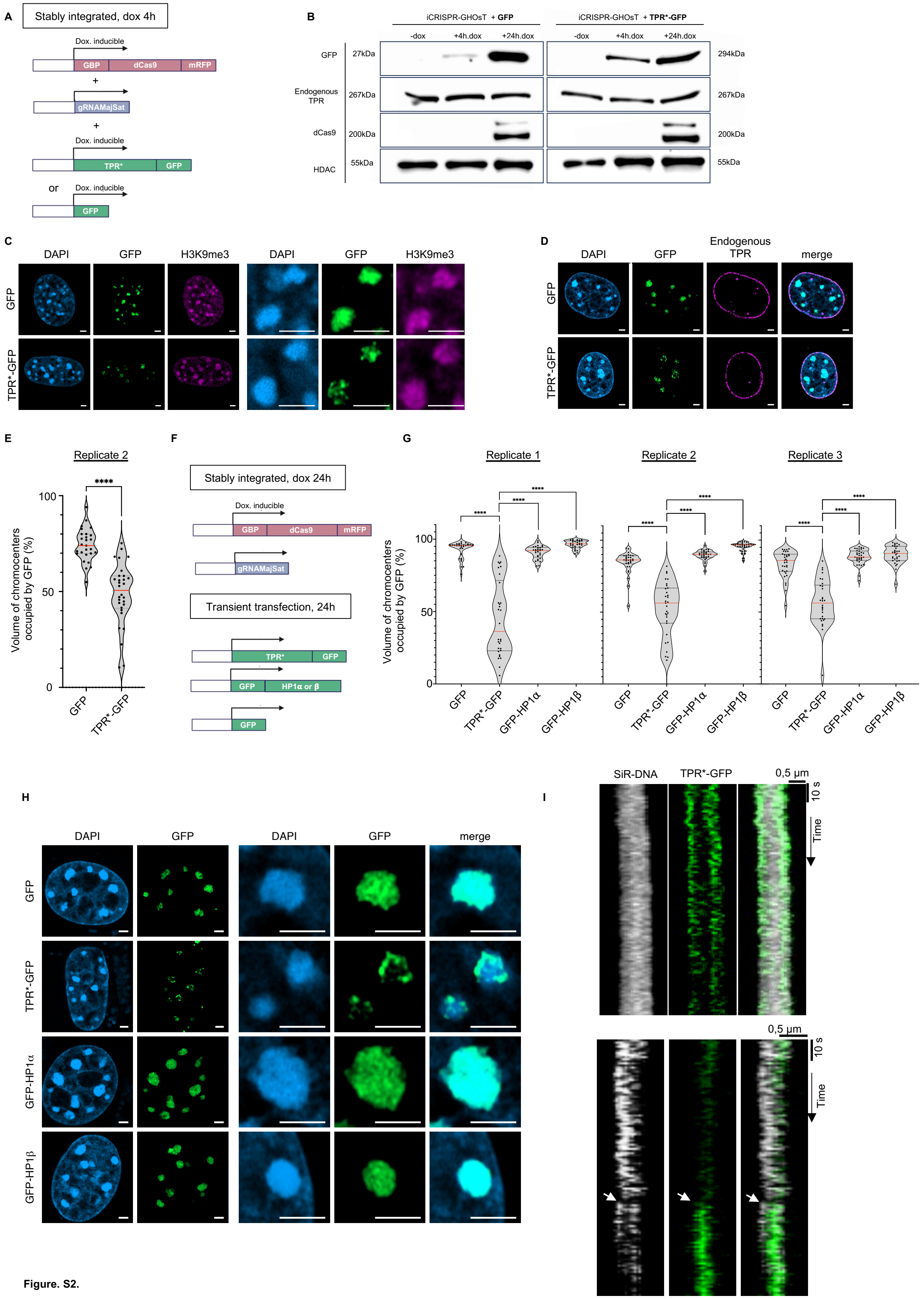
A
Stably integrated, dox 4h
Dox. inducible
GBP
dCas9
mRFP
+
gRNAMajSat
+
Dox. inducible
TPR*
GFP
or
Dox. inducible
GFP
B
iCRISPR-GHOsT + GFP
iCRISPR-GHOsT + TPR*-GFP
-dox
+4h.dox
+24h.dox
-dox
+4h.dox
+24h.dox
GFP
27kDa
294kDa
Endogenous TPR
267kDa
267kDa
dCas9
200kDa
200kDa
HDAC
55kDa
55kDa
C
DAPI
GFP
H3K9me3
DAPI
GFP
H3K9me3
GFP
TPR*-GFP
D
DAPI
GFP
Endogenous TPR
merge
GFP
TPR*-GFP
E
Replicate 2
****
100
50
0
Volume of chromocenters occupied by GFP (%)
GFP
TPR*-GFP
F
Stably integrated, dox 24h
Dox. inducible
GBP
dCas9
mRFP
gRNAMajSat
Transient transfection, 24h
TPR*
GFP
GFP
HP1α or β
GFP
G
Replicate 1
Replicate 2
Replicate 3
****
100
50
0
Volume of chromocenters occupied by GFP (%)
GFP
TPR*-GFP
GFP-HP1α
GFP-HP1β
H
DAPI
GFP
DAPI
GFP
merge
GFP
TPR*-GFP
GFP-HP1α
GFP-HP1β
I
SiR-DNA
TPR*-GFP
0,5 µm
10 s
Time
0,5 µm
10 s
Time

Figure. S2.

25

**Fig. S2. TPR* spatially segregates from heterochromatin.**

(A) Schematic of constructs expressed in iCRISPR-GHOsT cell lines in which doxycycline-inducible GFP or TPR*-GFP constructs have been stably integrated. (B) Immunoblots of protein extracts from iCRISPR-GHOsT cell lines in which doxycycline-inducible GFP or TPR*-GFP constructs have been stably integrated, without, 4h and 24h upon doxycyclin induction. (C) Airyscan pictures of immunofluorescence against H3K9me3 (purple) in iCRISPR-GHOsT cell lines in which doxycycline-inducible GFP or TPR*-GFP (green) constructs have been stably integrated, 4h upon doxycycline induction. DAPI is shown in blue. Right panel shows a zoom on two chromocenters for each condition. (D) Airyscan pictures of immunofluorescence against the endogenous TPR protein (purple) in iCRISPR-GHOsT cell lines in which doxycycline-inducible GFP or TPR*-GFP (green) constructs have been stably integrated, 4h upon doxycycline induction. DAPI is shown in blue. The TPR antibody used for this immunofluorescence does not recognize TPR-GFP or TPR*-GFP constructs. (E) Volume of chromocenters occupied by GFP signal (%) after targeting of GFP or TPR*-GFP to chromocenters in iCRISPR-GHOsT cell lines in which doxycycline-inducible GFP or TPR*-GFP constructs have been stably integrated, 4h upon doxycycline induction. A minimum of 30 cells per condition has been analyzed. Each point corresponds to one cell. The red line indicates the median. One replicate is shown here. Significance was calculated with Mann-Whitney test (****$p<0.0001$). (F) Schematic of constructs expressed upon transfection of GFP, TPR*-GFP or GFP-HP1 constructs in iCRISPR-GHOsT cells. (G) Volume of chromocenters occupied by GFP signal (%) after targeting of GFP or TPR*-GFP to chromocenters upon transfection of GFP, TPR*-GFP or GFP-HP1 constructs in iCRISPR-GHOsT cells, 24h upon transfection and doxycycline induction. A minimum of 30 cells per condition has been analyzed. Each point corresponds to one cell. The red line indicates the median. Three independent biological replicates are shown here. Significance was calculated with Mann-Whitney test (****$p<0.0001$). (H) Airyscan pictures of DAPI (blue), GFP, TPR*-GFP or GFP-HP1 (green) upon their transfection in iCRSIPR-GHOsT cells, 24h post-transfection and doxycycline induction. (I) Example kymographs generated along a line crossing the center of a chromocenter. The white arrow highlights rupture of heterochromatin domain upon TPR*-GFP condensate growth.

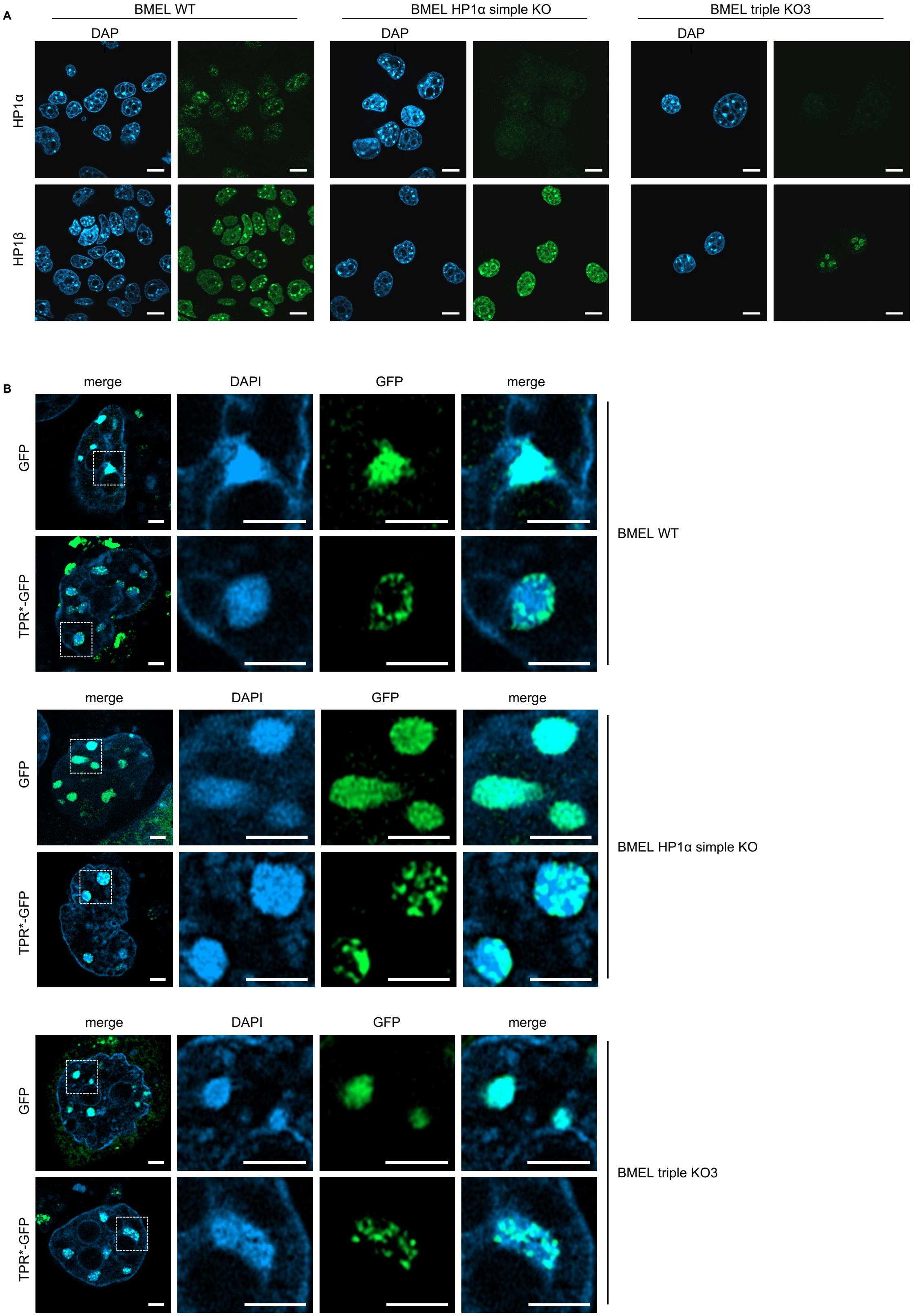
A
BMEL WT
BMEL HP1α simple KO
BMEL triple KO3
DAP
DAP
DAP
HP1α
HP1β
B
merge
DAPI
GFP
merge
GFP
TPR*-GFP
BMEL WT
merge
DAPI
GFP
merge
GFP
TPR*-GFP
BMEL HP1α simple KO
merge
DAPI
GFP
merge
GFP
TPR*-GFP
BMEL triple KO3

58

**Figure. S3.**

**Fig S3. HP1 proteins are not necessary for TPR* spatial segregation from heterochromatin.**

(A) Airyscan pictures of immunofluorescence against HP1$\alpha$ or $\beta$ in WT, HP1$\alpha$ or HP1 triple KO BMEL cells (green). DAPI is shown in blue. (B) Airyscan pictures of DAPI (blue), GFP or TPR*-GFP (green) upon their co-transfection with MajSat gRNAs and GBP-dCAS9-mRFP in WT, HP1$\alpha$ or HP1 triple KO BMEL cells, 24h post-transfection.

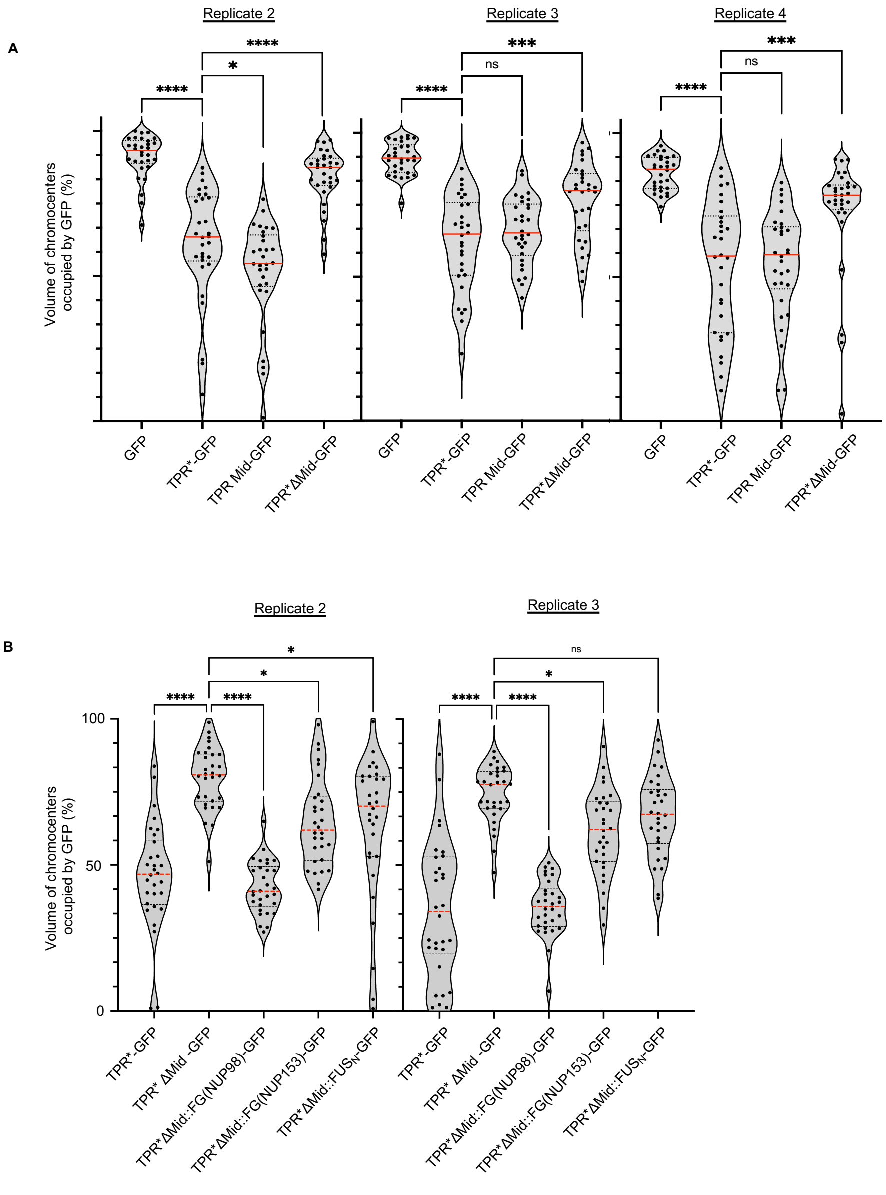
A
Replicate 2
Replicate 3
Replicate 4
Volume of chromocenters occupied by GFP (%)
****
*
****
***
ns
GFP
TPR*-GFP
TPR Mid-GFP
TPR*ΔMid-GFP
B
Replicate 2
Replicate 3
100
50
0
TPR*-GFP
TPR* ΔMid -GFP
TPR*ΔMid::FG(NUP98)-GFP
TPR*ΔMid::FG(NUP153)-GFP
TPR*ΔMid::FUSN-GFP

Figure. S4.

66

**Fig. S4. TPR biomolecular condensation is necessary and sufficient for spatial segregation from heterochromatin.**

(A) Volume of chromocenters occupied by GFP signal (%), after targeting GFP, TPR*-GFP and TPR*-GFP mutants to chromocenters upon their co-transfection with MajSat gRNAs in NIH3T3 cells in which a doxycycline-inducible GBP-dCas9-mRFP construct has been stably integrated, 24h post-transfection and doxycycline induction. A minimum of 30 cells per condition has been analyzed. Each point corresponds to one cell. The red line indicates the median. Three replicates are shown on this graph. Significance was calculated with Mann-Whitney test (ns: non-significant, *p<0.05 ***p<0.001, ****p<0.0001). (B) Volume of chromocenters occupied by GFP signal (%), after targeting TPR*-GFP or TPR* chimeras-GFP to chromocenters after their transfection in iCRISPR-GHOsT cell lines, 24h upon transfection and doxycyclin induction. A minimum of 30 cells per condition has been analyzed. Each point corresponds to one cell. The red line indicates the median. Two replicates are shown. Significance was calculated with Mann-Whitney test (ns: non-significant, *p<0.05, ****p<0.0001).

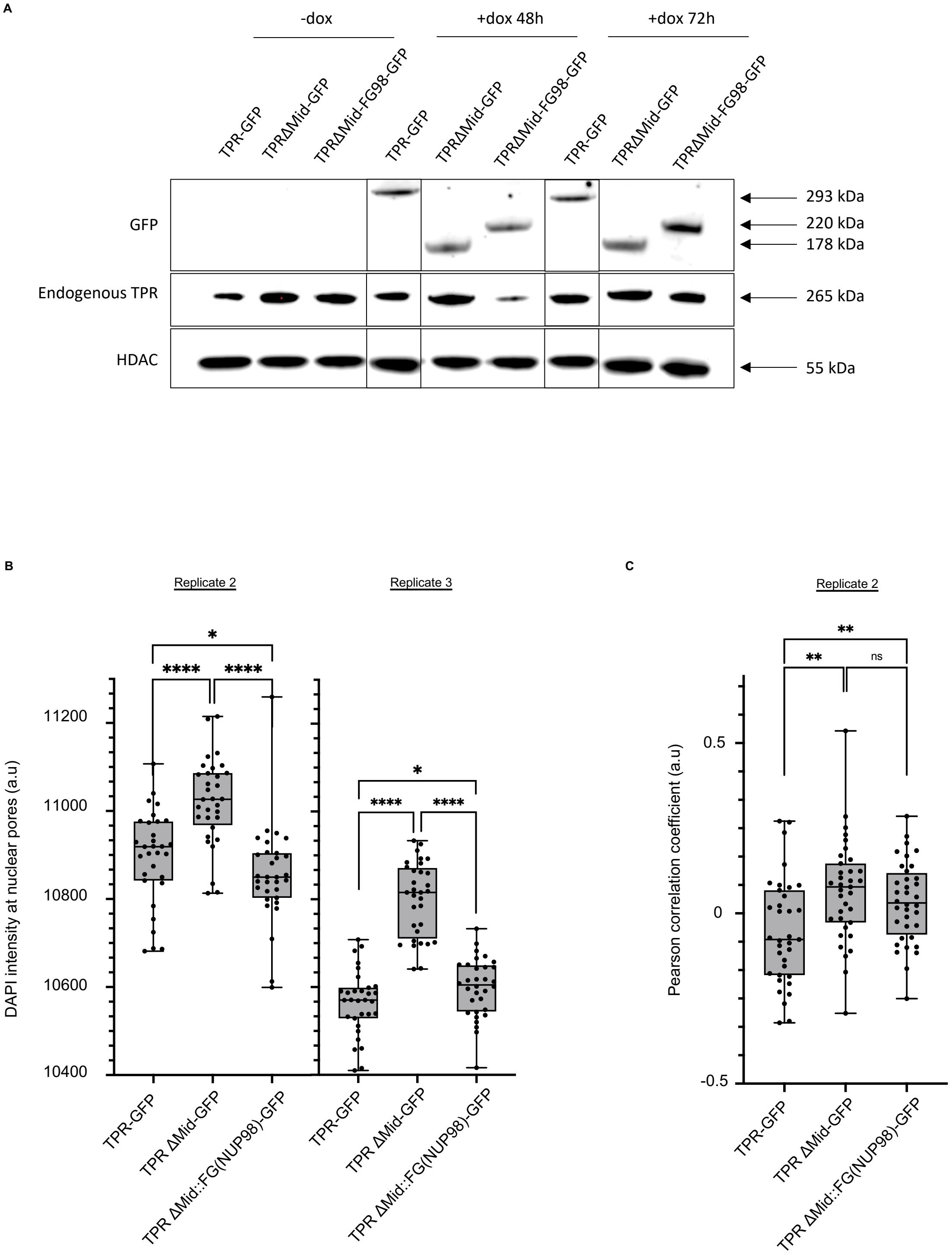
A
-dox
+dox 48h
+dox 72h
TPR-GFP
TPRΔMid-GFP
TPRΔMid-FG98-GFP
GFP
Endogenous TPR
HDAC
293 kDa
220 kDa
178 kDa
265 kDa
55 kDa
B
Replicate 2
Replicate 3
DAPI intensity at nuclear pores (a.u)
11200
11000
10800
10600
10400
*
****
TPR-GFP
TPR ΔMid-GFP
TPR ΔMid::FG(NUP98)-GFP
C
Replicate 2
Pearson correlation coefficient (a.u)
0.5
0
-0.5
**
ns

83 **Figure. S5.**

**Fig. S5. TPR molecular dynamics mediated by its disordered coiled coil is necessary and sufficient for heterochromatin exclusion at nuclear pores.**

(A) Immunoblots of protein extracts from NIH3T3 cells stably expressing either TPR-GFP, TPRΔMid-GFP or TPRΔMid::FG(Nup98)-GFP chimera under the control of a doxycycline inducible promoter, without, 48h and 72h upon doxycyclin induction. (B) DAPI intensity at nuclear pores in TPR-GFP, TPRΔMid-GFP or TPRΔMid::FG(Nup98)-GFP expressing cells, as in (A), 48h upon doxycycline induction. NUP153 immunofluorescence staining was used to segment nuclear pores in 3D. A minimum of 30 cells per condition has been analyzed. The interquartile range (IQR) is depicted by the box with the median represented by the center line. Whiskers maximally extend to 1.5 × IQR. Two replicates are shown here. Significance was calculated with Mann-Whitney test (*p<0.05, ****p<0.0001). (D) Pearson correlation coefficient between GFP intensity and DAPI intensity along the nuclear envelope in TPR-GFP, TPRΔMid-GFP or TPRΔMid::FG(Nup98)-GFP expressing cells, as in (A), 48h upon doxycycline addition. The interquartile range (IQR) is depicted by the box with the median represented by the center line. Whiskers maximally extend to 1.5 × IQR. A minimum of 30 cells per condition has been analyzed. One replicate is shown here. Significance was calculated with Mann-Whitney test (ns: non-significant, **p<0.01).

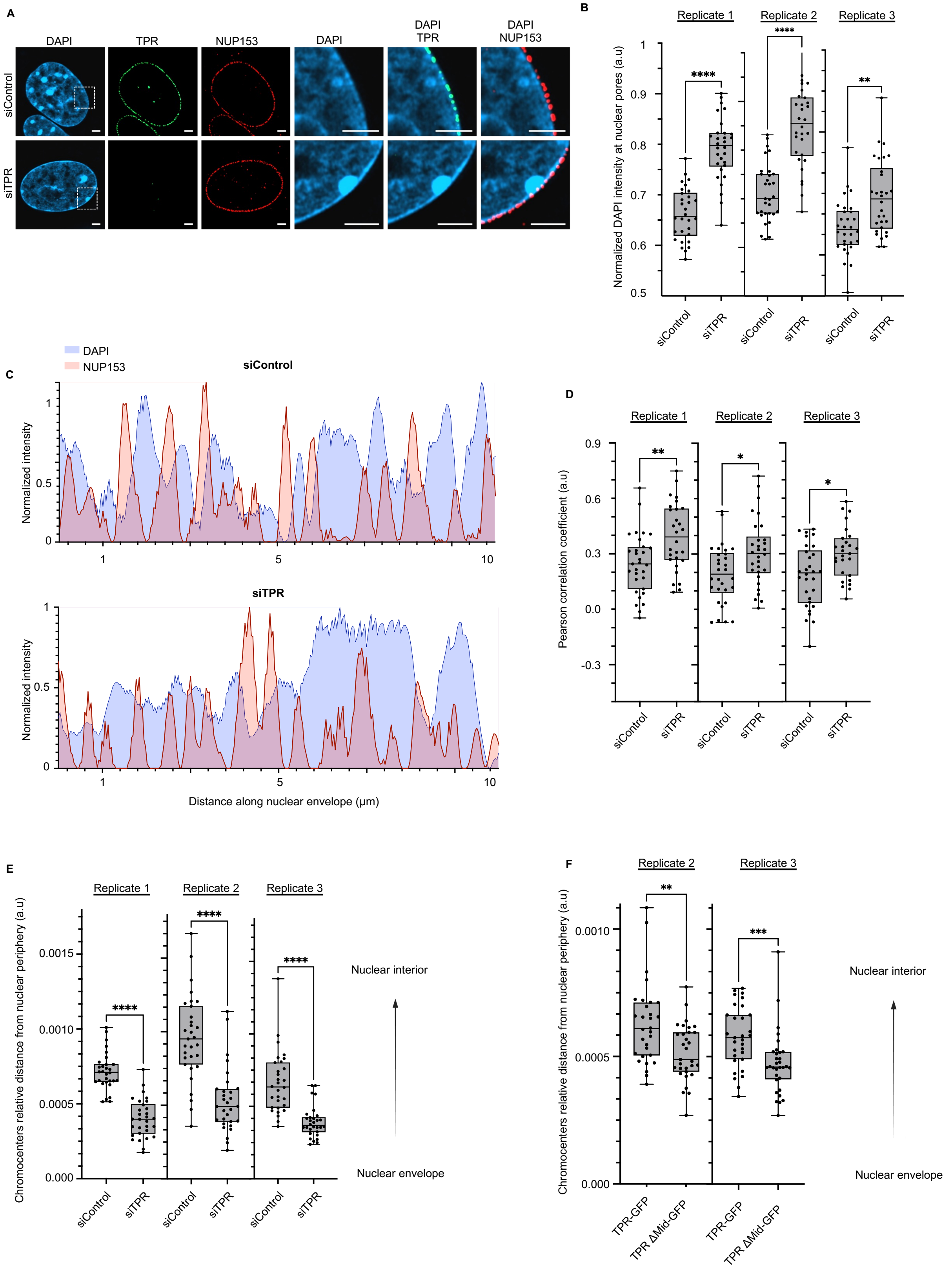


103 Figure. S6.

**Fig. S6. TPR is necessary for the maintenance of chromatin organization.**

(A) Airyscan pictures of Immunofluorescence against TPR (green) and NUP153 (red) in NIH3T3 cells treated with non-targeting siRNAs (siControl) or siRNAs against TPR (siTPR). DAPI is shown in blue. A single stack is shown here. (B) DAPI intensity at nuclear pores in NIH3T3 cells treated with siControl or siTPR, as in (A). NUP153 immunofluorescence staining was used to segment nuclear pores. A minimum of 30 cells per condition has been analyzed. The interquartile range (IQR) is depicted by the box with the median represented by the center line. Whiskers maximally extend to 1.5 × IQR. Three independent biological replicates are shown here. Significance was calculated with Mann-Whitney test (**p<0.01, ****p<0.0001). (C) Intensity of DAPI and NUP153 signals along the nuclear envelope of one representative cell treated with siControl or siTPR as in (A). (D) Pearson correlation coefficient between NUP153 staining intensity and DAPI intensity along the nuclear envelope in NIH3T3 cells treated with siControl or siTPR, as in (A). The interquartile range (IQR) is depicted by the box with the median represented by the center line. Whiskers maximally extend to 1.5 × IQR. A minimum of 30 cells per condition has been analyzed. Three independent biological replicates are shown here. Significance was calculated with Mann-Whitney test (**p<0.01, *p<0.05). (E) Chromocenter distance to the nuclear periphery normalized to nuclear volume in NIH3T3 cells treated with siControl or siTPR, as in (A). A minimum of 30 cells per condition has been analyzed. The interquartile range (IQR) is depicted by the box with the median represented by the center line. Whiskers maximally extend to 1.5 × IQR. Three independent biological replicates are shown here. Significance was calculated with Mann-Whitney test (****p<0.0001). (F) Chromocenter distance to the nuclear periphery normalized to nuclear volume in NIH3T3 cells stably expressing either TPR-GFP or TPRΔMid-GFP under the control of a doxycycline inducible promoter, 48h upon doxycycline addition. A minimum of 30 cells per condition has been analyzed. The interquartile range (IQR) is depicted by the box with the median represented by the center line. Whiskers maximally extend to 1.5 × IQR. Two independent biological replicates are shown here. Significance was calculated with Mann-Whitney test (**p<0.01, ***p<0.001).

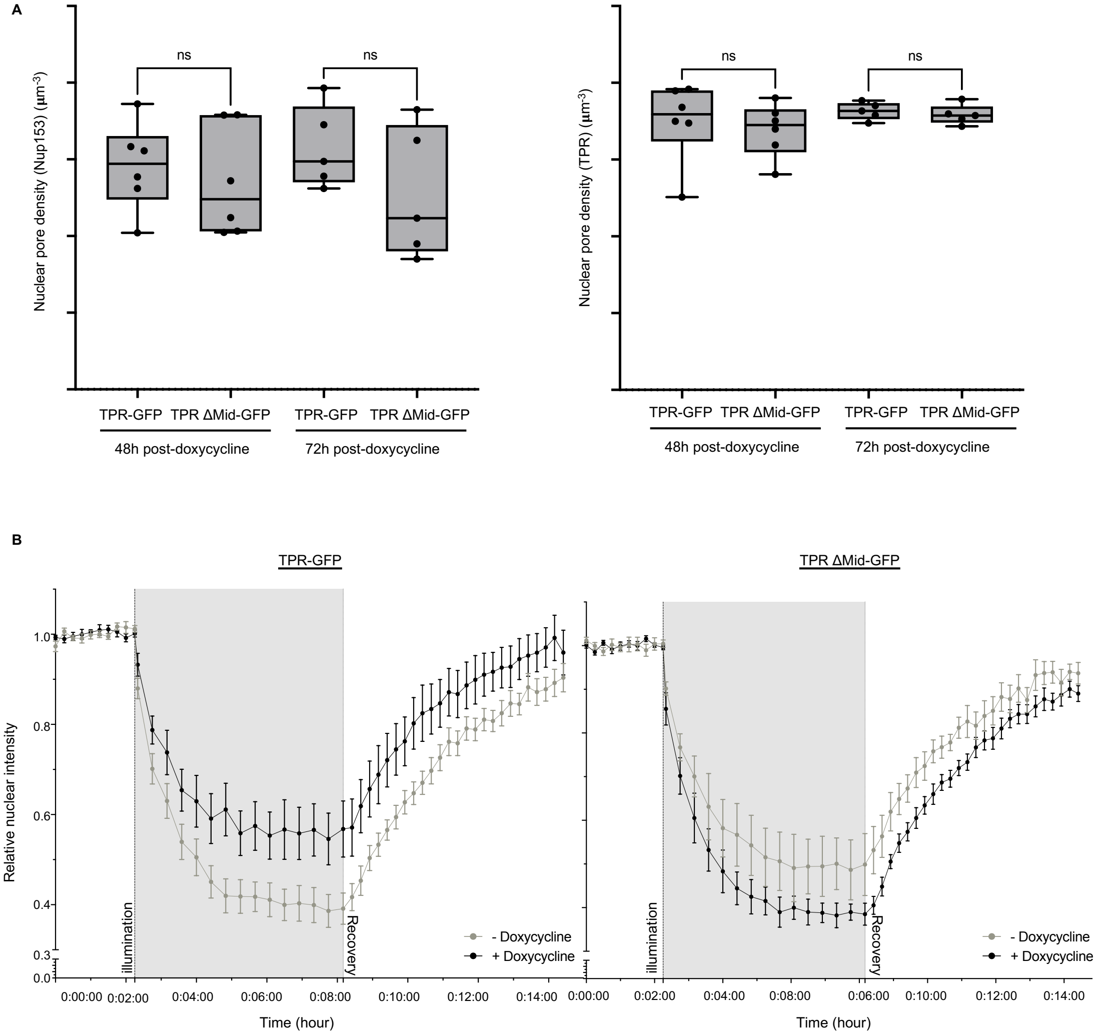


134 **Figure. S7.**

**Fig S7. TPR disordered coiled coil defines a new permeability barrier at the nuclear pore basket.**

(A) Nuclear pore density (total nuclear pore number normalized to nuclear volume) in NIH3T3 cells stably expressing either TPR-GFP or TPRΔMid-GFP under the control of a doxycycline inducible promoter, 48h or 72h upon doxycycline addition. Nuclear pores have been segmented in 3D using either NUP153 immunostaining (left panel) or TPR immunostaining (right panel). Each dot represents the mean of nuclear pore density measured in a minimum of 30 cells per condition. Six independent biological replicates are shown as individual dots here. The interquartile range (IQR) is depicted by the box with the median represented by the center line. Whiskers maximally extend to 1.5 × IQR. Significance was calculated with Anova test (n.s.: non-significant). (B) Quantification of the nuclear intensity of the light-inducible probe LEXY, normalized to the mean intensity obtained in the pre-illumination phase in NIH3T3 expressing (+doxycycline 72h) or not (-doxycycline) either TPR-GFP (left panel) or TPRΔMid-GFP (right panel). Upon 458 nm laser illumination (t = 2.25 min), LEXY translocates from the nucleus to the cytoplasm. After stopping the laser illumination (t = 8.10 min), LEXY is progressively imported to its initial nuclear localisation. One replicate out of the three is shown as an example, a minimum of 5 cells were analysed per replicate per condition.

**Movie S1. TPR* forms biomolecular condensates in transfected cells.** Time lapse imaging (1 image/10 seconds) 15h upon transfection of TPR*-GFP in NIH3T3 cells.

**Movie S2. TPR* forms biomolecular condensates early upon the expression of TPR*-GFP construct.** Time lapse imaging (1 image/10 seconds) of TPR*-GFP expressing NIH3T3 cells, 4h upon doxycycline induction.

**Movie S3.** Time lapse imaging (one image/second) of TPR*-GFP (green) targeted to chromocenters (marked with SiR-DNA, grey) in iCRISPR-GHOsT cells, 4h upon doxycycline induction.

**Table S1- Plasmids used and generated in this study**

| Plasmids | Sources/Reference | Mammalian Antibiotics Resistance | Promotor |
|---|---|---|---|
| GBP-dCas9-mRFP | Gift from S. Bultmann doi: 10.1080/19491034.2017.1292194 | - | CMV enhancer - Chicken β Promotor- chimeric intron |
| gRNA_MajSat | Gift from S. Polo doi: 10.1038/s41467-021-22575-5 | - | U6 promotor |
| GFP | This Paper | - | CMV enhancer - Chicken β Promotor - chimeric intron |
| TPR*-GFP (L458D & M489D) | This Paper, cloned from vector gifted by V. C. Cordes Doi:10.1091/mbc.12.8.2433 | - | CMV enhancer - Chicken β Promotor - chimeric intron |
| TPR*Nter-GFP | This Paper | - | CMV IE94 promoter |
| TPR Mid-GFP | This Paper | - | CMV IE94 promoter |
| TPR Cter-GFP | This Paper | - | CMV IE94 promoter |
| TPRΔNter-GFP | This Paper | - | CMV IE94 promoter |
| TPRΔMid-GFP | This Paper | - | CMV IE94 promoter |
| TPR*ΔMid-GFP | This Paper | - | CMV IE94 promoter |
| TPR*ΔCter-GFP | This Paper | - | CMV IE94 promoter |
| GFP- Nup153 | Addgene #64268 | - | CMV enhancer – CMV promotor |
| GFP-Nup98 | Gift from Maureen A. Powers doi: 10.1091/mbc.01-11-0538. | - | CMV enhancer – CMV promotor |
| $FUS_N$-GFP | This Paper, cloned from vector in doi : 10.1016/j.cell.2018.10.057. | - | CMV IE94 promoter |
| FGsNup98-GFP | This Paper | - | CMV IE94 promoter |
| FGsNup153-GFP | This Paper | - | CMV IE94 promoter |
| TPR*ΔMid::FG(Nup98)-GFP | This Paper | - | CMV IE94 promoter |

| | | | |
|---|---|---|---|
| TPR*ΔMid::FG(Nup153)-GFP | This Paper | - | CMV IE94 promoter |
| TPR*ΔMid::FUS$_N$-GFP | This Paper | - | CMV IE94 promoter |
| Lenti_TetON_GBP-dCas9-mRFP | This Paper | Neomycin | TetON-TetR |
| pKLV2.3_gRNAMajSat | This Paper | Hygromycin | - |
| Lenti_TetON_GFP | This Paper | Puromycin | TetON-TetR |
| Lenti_TetON_Piggybac_TPR*-GFP | This Paper | Puromycin | TetON-TetR |
| Lenti_TetON_TPR-GFP | This Paper | Puromycin | TetON-TetR |
| Lenti_TetON_TPRΔMid-GFP | This Paper | Puromycin | TetON-TetR |
| Lenti_TetON_TPRΔMid:FGsNup98-GFP | This Paper | Puromycin | TetON-TetR |
| HIV-1 gag/pol (psPAX) | Addgene #12260 | - | CMV enhancer - Chicken β Promotor - chimeric intron |
| VSV-G (pMD2.G) | Addgene #12259 | - | CMV enhancer – CMV promotor |
| Transposase | Giacomo Cavalli | - | CMV enhancer – CMV promotor |

**Table S2- Cell lines used and generated in this study**

| Stable cell lines | Parental cell line | Inducible Expression | Antibiotic Resistance |
|---|---|---|---|
| NIH3T3 (ATCC® CRL-1658™) | - | no | - |
| HEK293T (ATCC® CRL-11268™) | - | no | - |
| HK_CRISPR_TPR-mEGFP (Cytion) | - | no | - |
| NIH3T3_GBP-dCas9-mRFP | NIH3T3 (ATCC® CRL-1658™) | Tet-ON (Dox-inducible) | Neomycin |
| iCRISPR GHoST : NIH3T3_GBP-dCas9-mRFP_gRNAMajSat | NIH3T3_GBP-dCas9-mRFP | Tet-ON (Dox-inducible) | Neomycin Hygromycin |
| NIH3T3_GBP-dCas9-mRFP_gRNAMajSat_GFP | NIH3T3_GBP-dCas9-mRFP_gRNAMajSat | Tet-ON (Dox-inducible) | Neomycin Hygromycin Puromycin |

| | | | |
|---|---|---|---|
| NIH3T3_GBP-dCas9-mRFP_gRNAMajSat_TPR*-GFP | NIH3T3_GBP-dCas9-mRFP_gRNAMajSat | Tet-ON (Dox-inducible) | Neomycin<br>Hygromycin<br>Puromycin |
| NIH3T3_TPR*-GFP | NIH3T3 | Tet-ON (Dox-inducible | Puromycin |
| NIH3T3_TPR-GFP | NIH3T3 | Tet-ON (Dox-inducible) | Puromycin |
| NIH3T3_TPRΔMid-GFP | NIH3T3 | Tet-ON (Dox-inducible) | Puromycin |
| NIH3T3_TPRΔMid::FG(Nup98)-GFP | NIH3T3 | Tet-ON (Dox-inducible) | Puromycin |
| BMEL WT | Doi:10.1038/s41388-020-1177-8 | | |
| BMEL HP1α KO | Doi:10.1038/s41388-020-1177-8 | | |
| BMEL TKO | Doi:10.1038/s41388-020-1177-8 | | |

**Table S3- Antibodies used in this study**

| Primary Antibodies | Usage | Concentration | Species | Sources/Reference |
|---|---|---|---|---|
| Anti-TPR | IF/WB | 1/500 | Rabbit | Abcam (ab84516) |
| Anti-Nup153 | IF | 1/200 | Mouse | Abcam (ab24700) |
| Anti-GFP | WB | 1/250 | Mouse | Merck #11814 |
| Anti-RFP | WB | 1/500 | Rabbit | Invitrogen #PA1-986 |
| Anti-HDAC1 | WB | 1/2000 | Rabbit | Invitrogen #PA1-860 |
| GFP booster anti-GFP AF488 | IF | 1/1000 | Alpaca | Fisher Scientific 17343333 |
| **Secondary Antibodies** | **Usage** | **Concentration** | **Species** | **Sources/Reference** |
| Anti-Rabbit AF488 | IF | 1/500 | Donkey | Lifetechnologies A-21206 |
| Anti-Rabbit AF647 | IF | 1/500 | Donkey | Lifetechnologies A-31573 |
| Anti-Mouse AF555 | IF | 1/500 | Donkey | Lifetechnologies A-31570 |
| Anti-Rabbit HRP | WB | 1/10.000 | Goat | Cell signaling no. 7074 |
| Anti-Mouse HRP | WB | 1/10.000 | Horse | Cell signaling no. 7076 |

**Table S4- Scripts and Imaris batches used for image analyses**

All the scripts and macros can be found at https://gitlab.com/boumendil1/narat-figueroa-2025.git

| Script/ Macro | Name |
|---|---|
| Script 1 | FunctionsS8_TPRchromo |
| Script 2 | Script_Analysis_Table_v8 |
| Script 3 | FunctionsS10_siTPR |
| Script 4 | Script_Analysis_Table_v10 |
| Script 5 | FunctionsS12_TPRpores |
| Script 6 | Script_Analysis_Table_v12 |
| IMARIS batch 1 | TPR_foci.icpx |
| Macro 1 | Fiji_macro.ijm |
| Script 7 | PearsonCoeff_siRNA |
| Script 8 | PearsonCoeff_TPRpores |
